# Inositol Pyrophosphate Dynamics Reveals Control of the Yeast Phosphate Starvation Program Through 1,5-IP_8_ and the SPX Domain of Pho81

**DOI:** 10.1101/2023.02.14.528555

**Authors:** Valentin Chabert, Geun-Don Kim, Danye Qiu, Guizhen Liu, Lydie Michaillat Mayer, K. Muhammed Jamsheer, Henning Jacob Jessen, Andreas Mayer

## Abstract

Eukaryotic cells control inorganic phosphate to balance its role as essential macronutrient with its negative bioenergetic impact on reactions liberating phosphate. Phosphate homeostasis depends on the conserved INPHORS signaling pathway that utilizes inositol pyrophosphates (IPPs) and SPX receptor domains. Since cells synthesize various IPPs and SPX domains bind them promiscuously, it is unclear whether a specific IPP regulates SPX domains in vivo, or whether multiple IPPs act as a pool. In contrast to previous models, which postulated that phosphate starvation is signaled by increased production of the IPP 1-IP_7_, we now show that the levels of all detectable IPPs of yeast, 1-IP_7_, 5-IP_7_ and 1,5-IP_8_, strongly decline upon phosphate starvation. Among these, specifically the decline of 1,5-IP_8_ triggers the transcriptional phosphate starvation response, the PHO pathway. 1,5-IP_8_ inactivates the cyclin-dependent kinase inhibitor Pho81 through its SPX domain. This stimulates the cyclin-dependent kinase Pho85-Pho80 to phosphorylate the transcription factor Pho4 and repress the PHO pathway. Combining our results with observations from other systems we propose a unified model where 1,5-IP_8_ signals cytosolic phosphate abundance to SPX proteins in fungi, plants, and mammals. Its absence triggers starvation responses.

**Significance statement:** Cytosolic P_i_ is of prime importance for cellular bioenergetics because P_i_ influences free energy of nucleotide hydrolysis and the metabolite fluxes through glycolysis and oxidative phosphorylation. Eukaryotic cells use the INPHORS pathway to signal P_i_ via SPX domains and their ligands, inositol pyrophosphates (IP_7_, IP_8_), which control P_i_ homeostasis through a network of target proteins that import, export, store or detoxify P_i_. Studies with different systems failed to yield a coherent model on this regulation.

We performed the first time-resolved profiling of the full isomer spectrum of inositol pyrophosphates in yeast and dissected the isomer that is relevant to intracellular P_i_ signaling. Our results can be combined with existing observations from plants, mammals, and other fungi to support a unified model of P_i_ signaling across all eukaryotic kingdoms, which is in accord with the fundamental importance of P_i_ management for metabolism.

## Introduction

Inorganic phosphate (P_i_) is an essential nutrient for all living systems. While required in large amounts for synthesis of nucleic acids, phospholipids and phosphorylated carbohydrates and proteins, an overaccumulation of P_i_ decreases the free energy provided by P_i_-liberating reactions, such as nucleotide hydrolysis, which might stall metabolism (1). In fungi, plants and animals, control of P_i_ homeostasis involves *myo*-inositol pyrophosphates (IPPs) and a family of evolutionarily conserved SPX domains, constituting the core of a postulated signaling pathway that we termed INPHORS (1, 2).

SPX domains interact with or form part of a large variety of proteins that affect P_i_ homeostasis by transporting P_i_ across membranes, converting it into polyphosphates (polyP) or other metabolites, or regulating P_i_-dependent transcription (3). Direct regulation of an SPX-containing protein by synthetic IPPs was shown for the polyP polymerase VTC, the P_i_ transporters Pho91 and the PHR transcription factors in plants (4–13). A firm link suggesting the SPX domain as the receptor for inositol pyrophosphate regulation was provided by point mutants in the SPX domain that rendered VTC either independent of activation by IPPs, or non-responsive to them (6). Structural analysis revealed that many of these mutations localized to a highly charged region that can bind inositol poly- and pyrophosphates with high affinity. However, this binding site discriminates poorly between different IPPs at the level of binding (6). By contrast, strong differences are observed in the agonist properties, leading to the suggestion that binding affinity is a poor predictor of IPP specificity and activity (1, 4, 12).

IPPs that can be found in a wide variety of organisms carry seven (IP_7_) or eight (IP_8_) phosphates occupying every position around myo-inositol ring. The IP_7_ isomers 1PP-InsP_5_ and 5PP-InsP_5_ carry diphosphate groups at the 1- or 5-position, respectively, and 1,5(PP)_2_-InsP_4_ carries two diphosphate groups at the 1 and 5 position. For convenience, we refer to these IPPs as 5-IP_7_, 1-IP_7_ and 1,5-IP_8_ from hereon. All three IPPs are linked to phosphate homeostasis by the fact that genetic ablation of the enzymes making them, such as IP6Ks (inositol hexakisphosphate kinases), PPIP5Ks (diphosphoinositol pentakisphosphate kinases) and ITPKs (inositol tris/tetrakisphosphate kinases), alters the phosphate starvation response. Considerable discrepancies exist in the assignment of these IPPs to different aspects of P_i_ homeostasis. In mammalian cells, 1,5-IP_8_ activates the only SPX-protein expressed in this system, the P_i_ exporter XPR1 (14, 15). 1,5-IP_8_ was also proposed as a regulator of the phosphate starvation response in *Arabidopsis* because 1,5-IP_8_, but not 5-IP_7_, promotes the interaction of SPX1 with the P_i_-responsive transcription factor PHR1 *in vitro* (10). On the other hand, quantitative measurements of the interaction of IPPs with rice SPX4 and its cognate P_i_-responsive transcription factor PHR2 revealed only a minor, twofold difference in the K_d_ values for 5-IP_7_ and 1,5-IP_8_ (7). 1-IP_7_, 5-IP_7_ and 1,5-IP_8_ all decrease upon P_i_ starvation and mutants lacking either of the enzymes necessary for their synthesis in plants, the ITPKs and the PPIP5Ks, induce the phosphate starvation response (10, 16–18). Analysis of respective *Arabidopsis* mutants revealed that P_i_ concentration in shoots correlates poorly with their content of IP_7_ and IP_8_ (16). While *itpk1* mutants, lacking one of the enzymes that generates 5-IP_7_, have similar 1,5-IP_8_ content as wildtype, their P_i_ content is almost two-fold increased and their 5-IP_7_ content is reduced by a factor of four. Inversely, mutants lacking the PPIP5K VIH2 show a 5-fold reduction of 1,5-IP_8_ but normal P_i_ content (16). Although 1,5-IP_8_ is the IPP that is the most responsive to Pi starvation and P_i_ re-feeding (16) this has rendered it difficult to clearly resolve whether IP_7_, IP_8_, or both, signal cellular P_i_ status to the P_i_ starvation program *in vivo*. That plants are composed of multiple source and sink tissues for phosphate may complicate the analysis, because systemic knockouts of IPP synthesizing enzymes may exert their main effect in a tissue that is different from the one where the starvation response is scored.

Pioneering studies on the regulation of the P_i_ starvation program of *S. cerevisiae*, the PHO pathway, concluded that this transcriptional response is triggered by an increase in 1-IP_7_ (19, 20), whereas subsequent studies in the pathogenic yeast *Cryptococcus neoformans* proposed 5-IP_7_ as the necessary signaling compound (21). Studies using the *S. cerevisiae* polyphosphate polymerase VTC as a model suggested that this enzyme, which is necessary for polyP accumulation under P_i_-replete conditions, is stimulated by 5-IP_7_ in vivo (4, 11, 22, 23), whereas studies in *S. pombe* proposed IP_8_ as the stimulator (24). *S. pombe* has similar enzymes for IP_8_ synthesis and hydrolysis as *S. cerevisiae* (25–29). In contrast to *S. cerevisiae*, genetic ablation of 1-IP_7_ and IP_8_ production in *S. pombe* leads to hyper-repression of the transcriptional phosphate starvation response (27, 30, 31) and interferes with the induction of the transcriptional phosphate starvation response. However, the transcriptional phosphate starvation response in *S. pombe* occurs through a different set of protein mediators. For example, it lacks homologs of Pho81 and uses Csk1 instead of Pho85-Pho80 for phosphate-dependent transcriptional regulation (32–35). Furthermore, PHO pathway promotors in *S. pombe* overlap with and are strongly regulated by lncRNA transcription units (36–38). It is hence difficult to compare the downstream events in this system to the PHO pathway of *S. cerevisiae*.

The discrepancies mentioned above could either reflect a true divergence in the signaling properties of different IPPs in different organisms, in which case a common and evolutionarily conserved signaling mechanism may not exist. Alternatively, the diverging interpretations could reflect limitations in the analytics of IPPs and in the *in vivo* assays for the fundamental processes of P_i_ homeostasis. The analysis of IPPs is indeed very challenging in numerous ways. Their cellular concentrations are very low, the molecules are highly charged, and they exist in multiple isomers that differ only in the positioning of the pyrophosphate groups. IPP analysis has traditionally been performed by ion exchange HPLC of extracts from cells radiolabeled through ^3^H-inositol (39). This approach requires constraining and slow labeling schemes, is costly and time-consuming. Furthermore, HPLC-based approaches in most cases did not resolve isomers of IP_7_ or IP_8_. These factors severely limited the number of samples and conditions that could be processed and the resolution of the experiments.

The recent use of capillary electrophoresis coupled to mass spectrometry (CE-MS) has dramatically improved the situation, permitting resolution of many regio-isomers of IP_7_ and IP_8_ without radiolabeling, and at superior sensitivity and throughput. We harnessed the potential of this method to dissect the role of the three known IPPs that accumulate in *S. cerevisiae*. We analyzed their impact on the PHO pathway, which is a paradigm for a P_i_-controlled transcriptional response and the regulation of phosphate homeostasis (1, 40, 41). Beyond this physiological function, however, the PHO pathway also gained widespread recognition as a model for promotor activation, transcription initiation, chromatin remodeling and nucleosome positioning (42). In the PHO pathway, the cyclin-dependent kinase inhibitor (CKI) Pho81 translates intracellular P_i_ availability into an activation of the cyclin-dependent kinase (CDK) complex Pho85-Pho80 (19, 43–45). At high P_i_, Pho85-Pho80 phosphorylates and inactivates the key transcription factor of the PHO pathway, Pho4, which then accumulates in the cytosol (46–48). During P_i_ starvation, Pho81 inhibits the Pho85-Pho80 kinase, leading to dephosphorylation and activation of Pho4 and the ensuing expression of P_i_-responsive genes (PHO genes).

The activation of Pho85-Pho80 through Pho81 has been explored in detail, leading to a series of highly influential studies that have gained wide acceptance in the field. A critical function was ascribed to 1-IP_7_ in activating the CDK inhibitor PHO81, allowing it to inhibit Pho85-Pho80 (19). IP_7_ concentration was reported to increase upon P_i_ starvation, and this increase was considered as necessary and sufficient to inactivate Pho85-Pho80 kinase through Pho81 and trigger the PHO pathway. The 1-IP_7_ binding site on Pho81 was mapped to a short stretch of 80 amino acids, the “minimum domain” (19, 20, 44, 49). This domain is in the central region and distinct from the N-terminal SPX domain of Pho81. Since overexpression of the minimum domain rescued P_i_-dependent regulation of the PHO pathway to some degree, the key regulatory function was ascribed to this minimum domain and the SPX domain was considered as of minor importance. By contrast, earlier studies based on unbiased mutagenesis, which subsequently received much less attention, had identified mutations in other regions of PHO81 with significant impact on PHO pathway activation (44, 50–53). The situation is complicated by further studies, which found global IP_7_ levels to decrease rather than increase upon P_i_ starvation (6, 14, 22, 54). The analytics used in most studies could not distinguish 1-IP_7_ and 5-IP_7_, however, leaving open the possibility that an increase of 1-IP_7_ might be masked by a decrease of a much larger pool of 5-IP_7_.

To resolve these discrepancies and revisit the regulation of the PHO pathway by IPPs and Pho81, we capitalized on recent advances in the non-radioactive analysis of inositol pyrophosphates through CE-MS, offering superior resolution, sensitivity, and throughput (55, 56). We combined comprehensive analyses of PHO pathway activation and IPP profiles in mutants of key enzymes involved in IPP metabolism of yeast to determine the IPP species relevant to PHO pathway control and their impact on Pho85-Pho80 kinase. This led us to a revised model of PHO pathway regulation.

## Results

So far, analysis of the role of IPPs in P_i_ homeostasis and P_i_ starvation responses relied mainly on the use of mutants which ablate one of the pathways of IPP synthesis (Fig. 1). Yet, as shown in plants, several IPPs can change in a similar manner upon P_i_ depletion or replenishment, and ablation of enzymes adding phosphate groups at the 1- or 5-positions of the inositol ring induces similar P_i_ starvation responses (10, 16). This renders it difficult to distinguish a role for an individual IPP from the alternative hypothesis that all IPPs collectively contribute to signaling. To dissect this issue in yeast, we performed time course analyses of mutants in IPP phosphatases and kinases, in which we correlate the levels of all IPPs with the induction of the phosphate starvation response in yeast. The rationale was to seek for upper or lower thresholds of IPP concentrations during PHO pathway induction and using those to sieve out the IPP responsible for signaling P_i_ starvation.

**Fig. 1:**
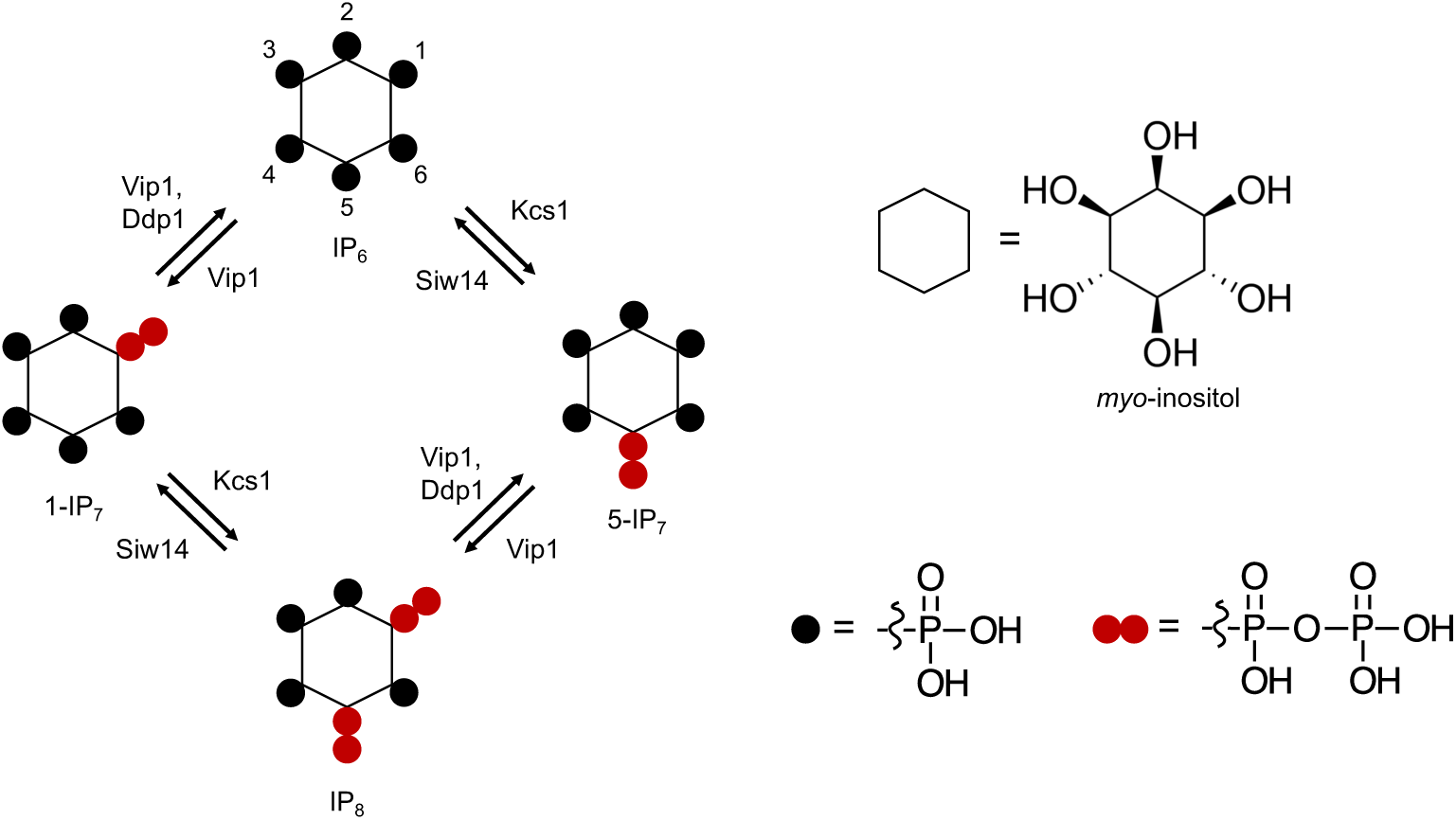
**Pathways of inositol pyrophosphate metabolism in *S. cerevisiae*.**

### Dynamics of the cytosolic concentrations of 5-IP_7_, 1-IP_7_ and 1,5-IP_8_

In yeast, the *myo*-inositol hexakisphosphate kinases Vip1 and Kcs1 generate 1-IP_7_ and 5-IP_7_, respectively, and they are both required for synthesis of 1,5-IP_8_ (Fig. 1) (57–61). Inositol pyrophosphatases, such as Ddp1 and Siw14 dephosphorylate these compounds at the 1- and 5-position, respectively (Fig. 1A) (22, 62, 63).

We analyzed the kinetics and the role of these inositol pyrophosphates in PHO pathway activation. To this end, yeasts were cultured in synthetic liquid (SC) media to early logarithmic phase and then transferred to P_i_-free SC medium. The cells were extracted with perchloric acid and inositol phosphates were analyzed by capillary electrophoresis coupled to electrospray ionization mass spectrometry (55). Three IPPs were detectable: 1-IP_7_, 5-IP_7_ and 1,5-IP_8_. We note that the CE-MS approach does not differentiate pyrophosphorylation of the inositol ring at the 1- and 3-positions. Thus, our assignments of the relevant species as 1-IP_7_ and 1,5-IP_8_ are based on previous characterization of the reaction products and specificities of IP6Ks and PPIPKs (57–61, 64).

Quantitation of IPPs by CE-MS was facilitated by spiking the samples with synthetic, ^13^C-labeled inositol pyrophosphate standards (65, 66). The recovery rate of inositol pyrophosphates during the extraction was determined by adding known quantities of synthetic standards to the cells already before the extractions. This demonstrated that 89 % of 1-IP_7_, 90 % of 5-IP_7_ and 75 % of 1,5-IP_8_ were recovered in the extract (Fig. S1). To estimate the cellular concentrations of these compounds, the volume of the cells was determined by fluorescence microscopy after staining of the cell wall with trypan blue (Fig. S2). This yielded an average cell volume of 42 fL. Detailed morphometric studies of yeast showed that the nucleus occupies around 8% of this volume and that all other organelles collectively account for approx. 18% (67). Taking this into account we can estimate the concentrations in the cytosolic space (including the nucleus, which is permeable to small molecules) of cells growing logarithmically on SC medium as 0.5 µM for 1-IP_7_, 0.7 µM for 5-IP_7_, 0.3 µM for 1,5-IP_8_ (Fig. 2A).

**Fig. 2.**
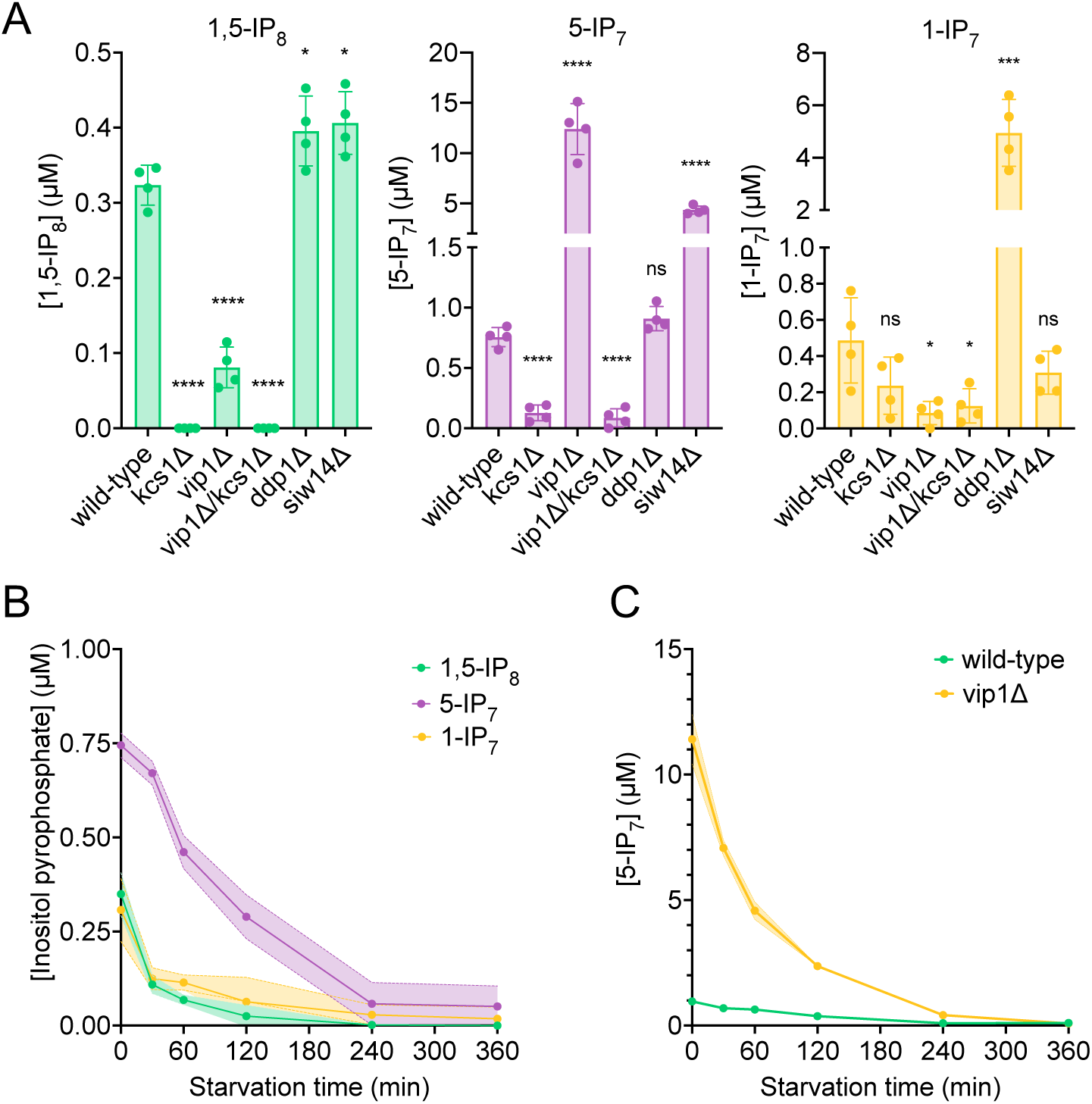
Cytosolic concentrations of 5-IP_7_, 1-IP_7_ and 1,5-IP_8_. (A) Inositol pyrophosphate concentrations in the cytosol. The indicated strains were grown logarithmically in SC medium containing 7.5 mM of P_i_ (30°C, 150 rpm, overnight). When OD_600nm_ reached 1 (1 × 10^7^ cells/ml), 1 ml of culture was extracted with perchloric acid and analyzed for IPPs by CE-ESI-MS. The y-axis provides the estimated cytosolic concentrations based on an average cell volume of 42 fL. Means (n=4) and standard deviations are indicated. **** p<0.0001; *** p<0.001; ** p<0.01; * p<0.05; n.s. not significant, tested with Student’s t-test. (B) Evolution of Inositol pyrophosphate species during P_i_ starvation. Cells were grown as in A, washed twice with P_i_ starvation medium and further incubated in P_i_ starvation medium. The inoculum for the samples bound to be extracted after different times of further incubation in starvation medium was adjusted such that all samples had similar OD_600nm_ at the time of harvesting (OD_600nm_=0.5 for 30 min and 60 min samples; OD_600nm_=0.4 for 120 min and 240 min samples; OD_600nm_=0.25 for 360 min samples). At the indicated times in starvation media, 1 ml aliquots were extracted and analyzed for IPPs as in A. The data was normalized by the number of cells harvested before calculating cytosolic concentrations. Means and standard deviations are given (n=3). (C) Depletion of 5-IP_7_ in starving *vip1Δ* cells. The indicated cells were grown in Pi-replete medium and then transferred to Pi starvation medium as in B. At the indicated times, samples were extracted and analyzed for 5-IP_7_ as in A. Means and standard deviations (n=4) are shown as solid lines and shaded areas, respectively.

Next, we determined the impact of Kcs1, Vip1, Siw14 and Ddp1 on the inositol pyrophosphate levels in the cells (Fig. 2A). 5-IP_7_ was not detected in the *kcs1Δ* mutant and 1-IP_7_ was strongly reduced in the *vip1Δ* strain. 1,5-IP_8_ was undetectable in *kcs1Δ* and decreased by 75% in *vip1Δ*. The nature of the residual 1,5-IP_8_ and 1-IP_7_ signals is currently unclear. They may represent IPPs synthesized by enzymes other than Kcs1 and Vip1, such as the inositol polyphosphate multi-kinases, which can also produce IPPs (16, 59, 68–70). Importantly, residual 1,5-IP_8_ and 1-IP_7_ were not observed in P_i_-starved wildtype cells (Fig. 2B). This may be due to presence of the Vip1 phosphatase activity, which is missing in *vip1Δ* cells, but which may quench weak production of IPPs such as 1-IP_7_ or 1,5-IP_8_ by other enzymes in wildtype cells. Since this aspect is not central to the question of our study it was not pursued further. *kcs1Δ* mutants showed a 2 to 3-fold decrease in 1-IP_7_, suggesting that the accumulation of 1-IP_7_ depends on 5-IP_7_. This might be explained by assuming that, in the wildtype, most 1-IP_7_ stems from the conversion of 5-IP_7_ to 1,5-IP_8_, followed by dephosphorylation of 1,5-IP_8_ to 1-IP_7_. A systematic analysis of this interdependency will require rapid pulse-labeling approaches for following the turnover of the phosphate groups, which are not yet established for inositol pyrophosphates (39, 65, 71, 72). An unexpected finding was the up to 20-fold overaccumulation of 5-IP_7_ in the *vip1Δ* mutant. By contrast, *ddp1Δ* cells showed normal levels of 5-IP_7_ and 1,5-IP_8_ but a 10-fold increase in 1-IP_7_. *siw14Δ* cells showed a 5-fold increase in 5-IP_7_, but similar levels of 1,5-IP_8_ and 1-IP_7_ as wildtype.

We performed time course experiments to analyze how IPP levels change under P_i_ withdrawal. 5-IP_7_ was the predominant inositol pyrophosphate species in wildtype cells growing on P_i_-replete media (Fig. 2B). Within 30 min of P_i_ starvation, the concentration of all three inositol pyrophosphate species rapidly decreased by 75% for 1,5-IP_8_, by 47% for 1-IP_7_, and by 40% for 5-IP_7_. This decline continued, so that 1-IP_7_ and 1,5-IP_8_ became undetectable and only 3% of 5-IP_7_ remained after 4 h, corresponding to a concentration below 50 nM. The high excess of 5-IP_7_ in *vip1Δ* cells also declined as soon as the cells were transferred to P_i_ starvation medium (Fig. 2C). After two hours of starvation, it was still two-fold above the concentration measured in P_i_-replete wildtype cells. Even after 3.5 to 4 h, P_i_-starved *vip1Δ* cells had just reached the 5-IP_7_ concentration of P_i_-replete wildtype cells. A comparable decline of all IPP species upon P_i_ starvation could be also observed in other fungi, such as *Schizosaccharomyces pombe* and *Cryptococcus neoformans* (Fig. S3), suggesting that this response is conserved.

Taken together, P_i_ starvation leads to a virtually complete depletion of all three inositol pyrophosphate species, with 1,5-IP_8_ declining faster than 1-IP_7_ and 5-IP_7_. Furthermore, inositol pyrophosphatase mutants provide the possibility to generate relatively selective increases in 5-IP_7_ and 1-IP_7_. We used this information to dissect the impact of 5-IP_7_, 1-IP_7_ and 1,5-IP_8_ on control of the PHO pathway.

### 1,5-IP8 signals cytosolic Pi levels to the PHO pathway

To this end, we correlated the measured inositol pyrophosphate concentrations to the induction of the PHO pathway. We assayed a key event of PHO pathway activation, partitioning of the fluorescently tagged transcription factor Pho4^yEGFP^ between the cytosol and the nucleus. Pho4 shuttles between nucleus and cytosol and its phosphorylation through Pho85-Pho80 favors Pho4 accumulation in the cytosol. The relocation of Pho4 can hence serve as an *in vivo* indicator of PHO pathway activation (73–75). It provides a readout for Pho85-Pho80 activity. *PHO4* was tagged at its genomic locus, making Pho4^yEGFP^ the sole source of this transcription factor. Pho4 relocation was assayed through automated image segmentation and analysis. The artificial intelligence-based segmentation algorithm recognized more than 90% of the cells in a bright-field image and delimited their nuclei based on a red-fluorescent nuclear mCherry marker (Fig. S4). This segmentation allows quantitative measurements of Pho4 distribution between the cytosol and nucleus in large numbers of cells. In addition, we assayed PHO pathway activation through fluorescent yEGFP reporters expressed from the *PHO5* (*prPHO5-yEGFP*) and *PHO84* (*prPHO84-yEGFP*) promotors. These are classical assays of PHO pathway activation, but their output is further downstream and hence comprises additional regulation, e. g. at the level of chromatin or RNA, or the direct activation of Pho4 through metabolites such as AICAR (76–80). Upon P_i_ withdrawal, both promotors are induced by the PHO pathway but the *PHO84* promotor reacts in a more sensitive manner and is induced more rapidly than the *PHO5* promotor (73).

In wildtype cells grown under P_i_-replete conditions, Pho4^yEGFP^ was cytosolic and the *PHO5* and *PHO84* promotors were inactive, indicating that the PHO pathway was repressed (Fig. 3). Within 30 min of starvation in P_i_-free medium, Pho4^yEGFP^ relocated into the nucleus and *PHO84* and *PHO5* were strongly induced. By contrast, *kcs1Δ* cells showed Pho4^yEGFP^ constitutively in the nucleus already under P_i_-replete conditions, and *PHO5* and *PHO84* promoters were activated. These cells have strongly reduced 1,5-IP_8_ and 5-IP_7_, and 50% less 1-IP_7_ than the wildtype, Thus, a decline of IPPs not only coincides with the induction of the P_i_ starvation program, but the genetic ablation of these compounds is sufficient for a forced triggering of this response in P_i_-replete conditions. Therefore, we explored the hypothesis that IPPs repress the PHO pathway, and that their loss upon P_i_ starvation creates the signal that activates the starvation response.

**Fig. 3.**
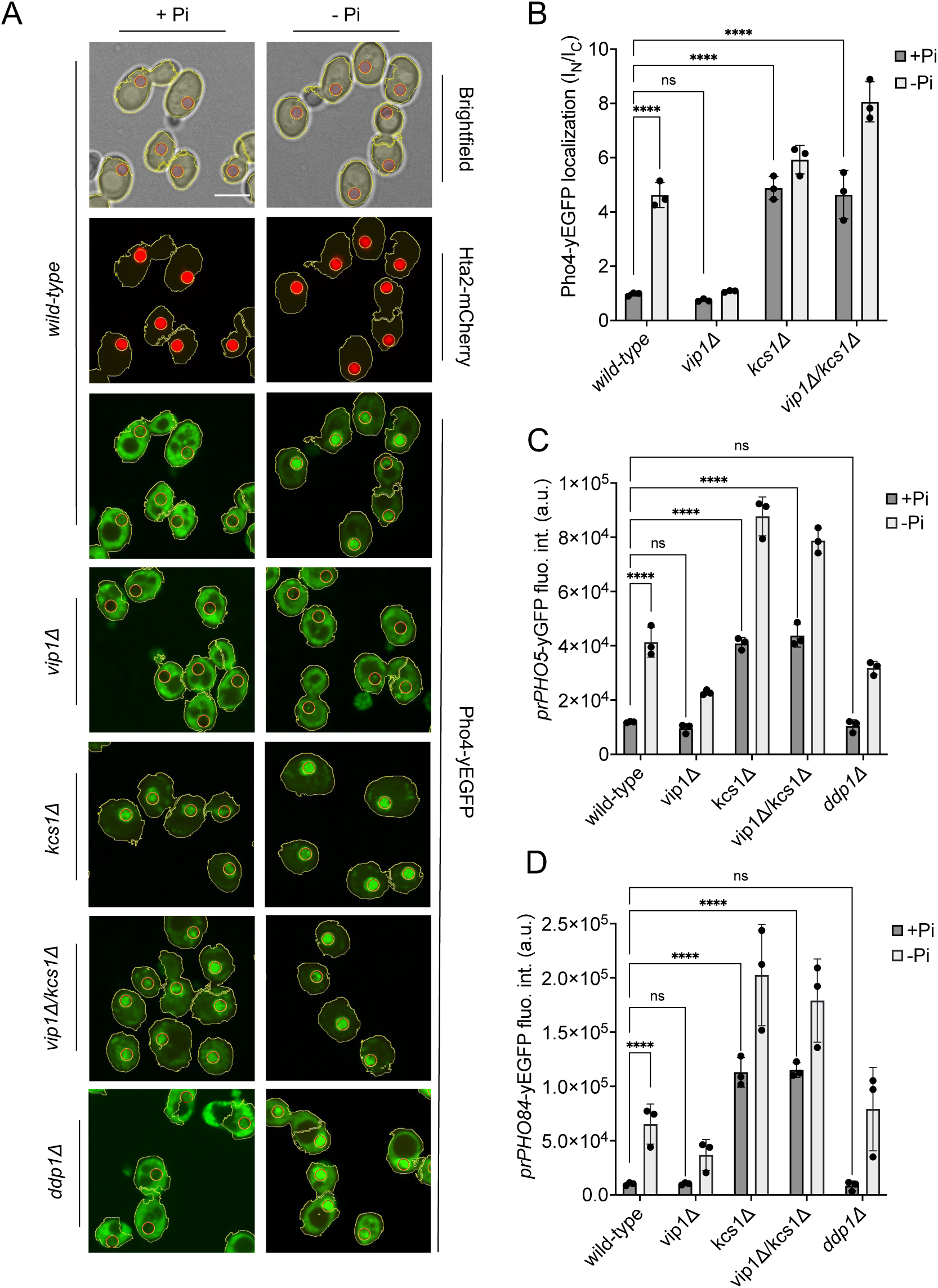
Inhibition of the PHO pathway by excessive 5-IP7. The indicated cells producing Pho4^yEGFP^ and the histone Hta2^mCherry^ as a nuclear marker were logarithmically grown in P_i_-replete SC medium, washed, and transferred to P_i_ starvation medium as in Fig. 2A. (A) Subcellular localization of Pho4^yEGFP^ was analyzed on a spinning disc microscope. Cells are shown in the presence of 7.5 mM of P_i_ (+ P_i_) or 30 min after the shift to P_i_ starvation (-P_i_) medium. Yellow lines surrounding the cells illustrate the segmentation performed by the algorithm that was used to quantify Pho4^yEGFP^ distribution in B. Scale bar: 5 μM. λ_ex_: 470 nm; λ_em_: 495-560 nm. (B) Average intensity of Pho4^yEGFP^ fluorescence was determined by automated image segmentation and analysis. Pho4^yEGFP^ localization is quantified by the ratio of the average fluorescence intensities in the nucleus over the average fluorescence intensity in the cytosol (I_N_/I_C_). 100 to 200 cells were analyzed per condition and experiment. n=3. Means and standard deviation are indicated. (C) Activation of the PHO5 promotor. Cells expressing the *prPHO5-yEGFP* reporter construct from a centromeric plasmid were grown in P_i_-replete medium (7.5 mM P_i_) as in Fig. 2A, and then shifted to P_i_ starvation medium or kept in P_i_ replete medium. After 4 h of further incubation, fluorescence intensity of the same number of cells was measured in a Spectramax EM microplate reader. λ_ex_: 480 nm; λ_em_: 510 nm. n=3. Means and standard deviations are indicated. (D) Activation of the PHO84 promotor. Cells expressing the *prPho84-yEGFP* reporter construct from a centromeric plasmid were treated and analyzed as in C. For B, C, and D: **** p<0.0001; *** p<0.001; ** p<0.01; * p<0.05.; n.s. not significant, tested with Turkey’s test.

In this case, we must explain the behavior of the *vip1Δ* mutation, which strongly reduces 1-IP_7_ and 1,5-IP_8_ and maintains the PHO pathway repressed in P_i_-replete medium. Upon withdrawal of P_i_, *vip1Δ* cells did not show nuclear relocation of Pho4 after 30 min and, even after 4h of starvation, *PHO5* was not expressed. The *PHO84* promoter remained partially repressed in comparison with the wildtype. These results are at first sight consistent with the proposal that 1-IP_7_ activates the PHO pathway (19, 20). Several further observations draw this hypothesis into question, however. First, all three inositol pyrophosphate species strongly decline upon P_i_ starvation instead of showing the increase postulated by Lee et al. Second, *vip1Δ* cells show a more than 15-fold overaccumulation of 5-IP_7_. Genetic ablation of this pool by deleting *KCS1* is epistatic to the *vip1Δ* mutation. A *kcs1Δ vip1Δ* double mutant constitutively activates the PHO pathway already in presence of P_i_, despite its strong reduction of 1-IP_7_ and the complete absence of 1,5-IP_8_. Third, *ddp1Δ* cells, which show a 10-fold quite selective increase in 1-IP_7_ under P_i_-replete conditions, did not induce the *PHO84* and *PHO5* promotors, nor did they show Pho4 accumulation in the nucleus in P_i_-replete medium. Thus, even a strong increase in 1-IP_7_ is not sufficient to activate the PHO pathway, while reductions of inositol pyrophosphates do this.

The data described above argues against a diminution of 1-IP_7_ as a critical factor for induction of the PHO pathway because *vip1Δ* cells, which have virtually no 1-IP_7_, do not constitutively activate the PHO pathway. By contrast, *vip1Δ kcs1Δ* double mutants, which not only have low levels of 1-IP_7_ and 1,5-IP_8_ but also strongly reduced 5-IP_7_, show constitutive PHO pathway activity (Fig. 3). We hence tested the hypothesis that a decline of 5-IP_7_ might trigger the PHO pathway. To define the critical concentration at which this happens in *vip1Δ* cells, we assayed the induction of the PHO pathway over time (Fig. 4). While Pho4^yEGFP^ was cytosolic in wild-type cells under high P_i_ supply, a large fraction of it rapidly relocated into the nucleus within the first 15 min of P_i_ starvation (Fig. 4 A,B). In *vip1Δ*, Pho4^yEGFP^ relocation followed a sigmoidal curve. The nuclear/cytosolic ratio strongly increased after 1 h, reaching a plateau after 3 h. Thus, relocation was stimulated when the 20-fold exaggerated 5-IP_7_ levels of the *vip1Δ* cells had declined below 5 µM (Fig. 2C), which is 6 to 7 times above the maximal concentration observed in P_i_-replete wildtype cells. Since 5-IP_7_ is the only IPP that exists in *vip1Δ* cells in significant amounts, this observation places the critical concentration of 5-IP_7_ that can repress the PHO pathway at 5 µM. In P_i_-replete wildtype cells, 5-IP_7_ never reaches 5 µM (Fig. 2) and, therefore, 5-IP_7_ is unlikely to be a physiological repressor of the PHO pathway in wildtype cells. The 5-IP_7_-dependent PHO pathway repression in *vip1Δ* cells must then be considered as a non-physiological reaction resulting from the strong overaccumulation of 5-IP_7_ in these cells.

**Fig. 4.**
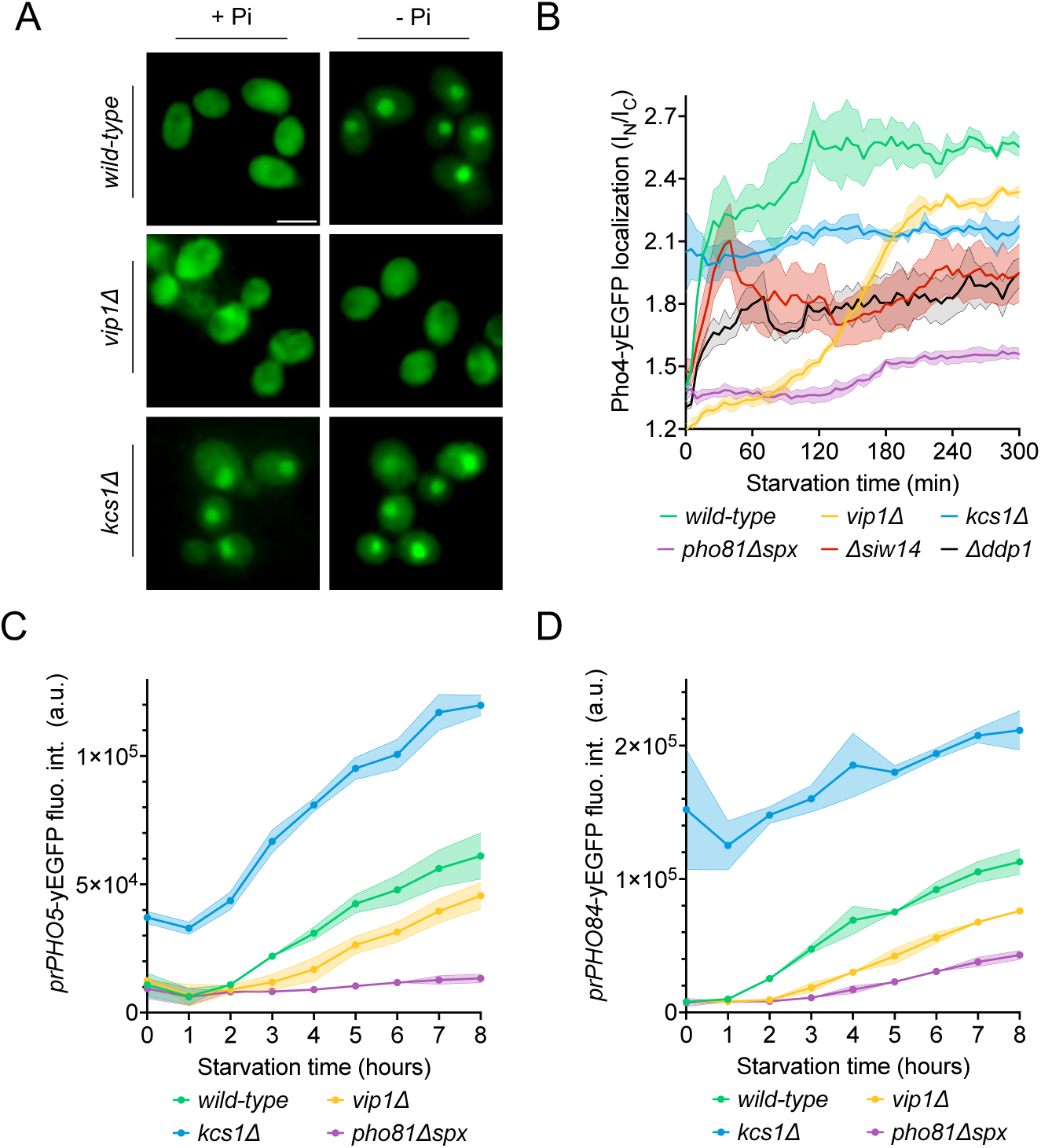
Effect of Vip1, Kcs1 and Pho81 on the time course of Pho4 translocation and PHO-gene activation. The indicated cells were logarithmically grown in P_i_-replete SC medium, washed, and transferred to P_i_ starvation medium as in Fig. 2A. At the indicated times after transfer to Pi-starvation medium, they were analyzed for Pho4 localization, *prPHO5-yEGFP* expression, or *prPHO84-yEGFP* expression, using assays as in Fig. 3. (A) Cells expressing Pho4^yEGFP^ and the histone Hta2^mCherry^ as a nuclear marker, shown in the presence of 7.5 mM of P_i_ (+ P_i_) or after 90 min of P_i_ starvation (-P_i_). Pictures were taken on a widefield fluorescence microscope equipped with a stage-top incubator (kept at 30°C) and an IBIDI flow chamber. Only the GFP-channel is shown. Scale bar = 5 μM. λ_ex_: 470 nm; λ_em_: 495-560 nm. (B) Distribution of Pho4^yEGFP^ between the nucleus and cytosol was quantified in the cells from A at various timepoints of P_i_ starvation. 100-200 cells were analyzed per condition and experiment at each timepoint. The solid lines and the shaded areas indicate the means and standard deviation, respectively. (C) Activation of the *PHO5* promotor. Cells expressing the *prPho5-yEGFP* reporter construct from a centromeric plasmid were grown in P_i_-replete medium, shifted to P_i_ starvation medium, and analyzed for GFP fluorescence intensity (as in Fig. 3C) at the indicated timepoints of starvation. λ_ex_: 480 nm; λ_em_: 510 nm. n=3. The solid lines and the shaded areas indicate the means and standard deviation, respectively. (D) Activation of the *PHO84* promotor. As in C, but with cells expressing *prPHO84-yEGFP* as reporter.

This interpretation is also consistent with the phenotype of *siw14Δ* cells. Although these cells accumulate 5-IP_7_, they accumulate it 4-fold less than *vip1Δ*, remaining below the 5 µM threshold where we should expect non-physiological repression of the PHO pathway. However, *siw14Δ* cells contain the same concentration of 1,5-IP_8_ as the wildtype (Fig. 2A), degrade it upon P_i_ starvation as the wildtype (Fig. S5), and relocalize Pho4^yEGFP^ with similar kinetics as the wildtype (Fig. 4B). Taken together with the evidence presented above, 1,5-IP_8_ thus ends up being the only IPP that correlates with the activation of the PHO pathway in all mutants and conditions. This points to 1,5-IP_8_ as the controller of the PHO pathway.

### The PHO pathway is controlled by the SPX domain of Pho81

Our results strongly argue against an increase of 1-IP_7_ as the key factor for inactivating the CDK Pho85-Pho80 through the CDK inhibitor Pho81. This prompted us to also re-evaluate the regulation of Pho81 because previous studies had proposed a small peptide from the central region of Pho81, called the minimum domain (amino acids 645-724), as a receptor for 1-IP_7_ that triggers the PHO pathway (19, 20). This model rests mainly on results with the isolated minimum domain used *in vitro* or, in highly overexpressed form, *in vivo*. This expression of the minimum domain outside its normal molecular context may be problematic because earlier work had suggested that the N-and C-terminal portions of Pho81, i. e. regions outside the minimum domain, could provide competing inhibitory and stimulatory functions for Pho81 (44).

We targeted the N-terminal SPX domain of Pho81 through various mutations, leaving the rest of Pho81 and its minimum domain intact (Fig. 5). First, we asked whether Pho81 lacking the N-terminal 200 amino acids, corresponding to a deletion of its SPX domain, could still activate the PHO pathway. This Pho81^ΔSPX^ is likely to be folded because a yEGFP-tagged version of Pho81^ΔSPX^ localized to the nucleus (Fig. 5C, D), which requires Pho81 to interact with nuclear import factors and with Pho85-Pho80 (49). We tested the effect of Pho81^ΔSPX^ on relocation of Pho4^yEGFP^ (Fig. 5 A, B). Whereas wild-type cells efficiently relocated Pho4^yEGFP^ to the nucleus upon P_i_ starvation, *pho81^ΔSPX^* cells partially maintained Pho4^yEGFP^ in the cytosol. Thus, while the SPX domain is not essential for nuclear targeting of Pho81, it is required for efficient nuclear relocation of Pho4 and the induction of the PHO pathway under P_i_ starvation.

**Fig. 5.**
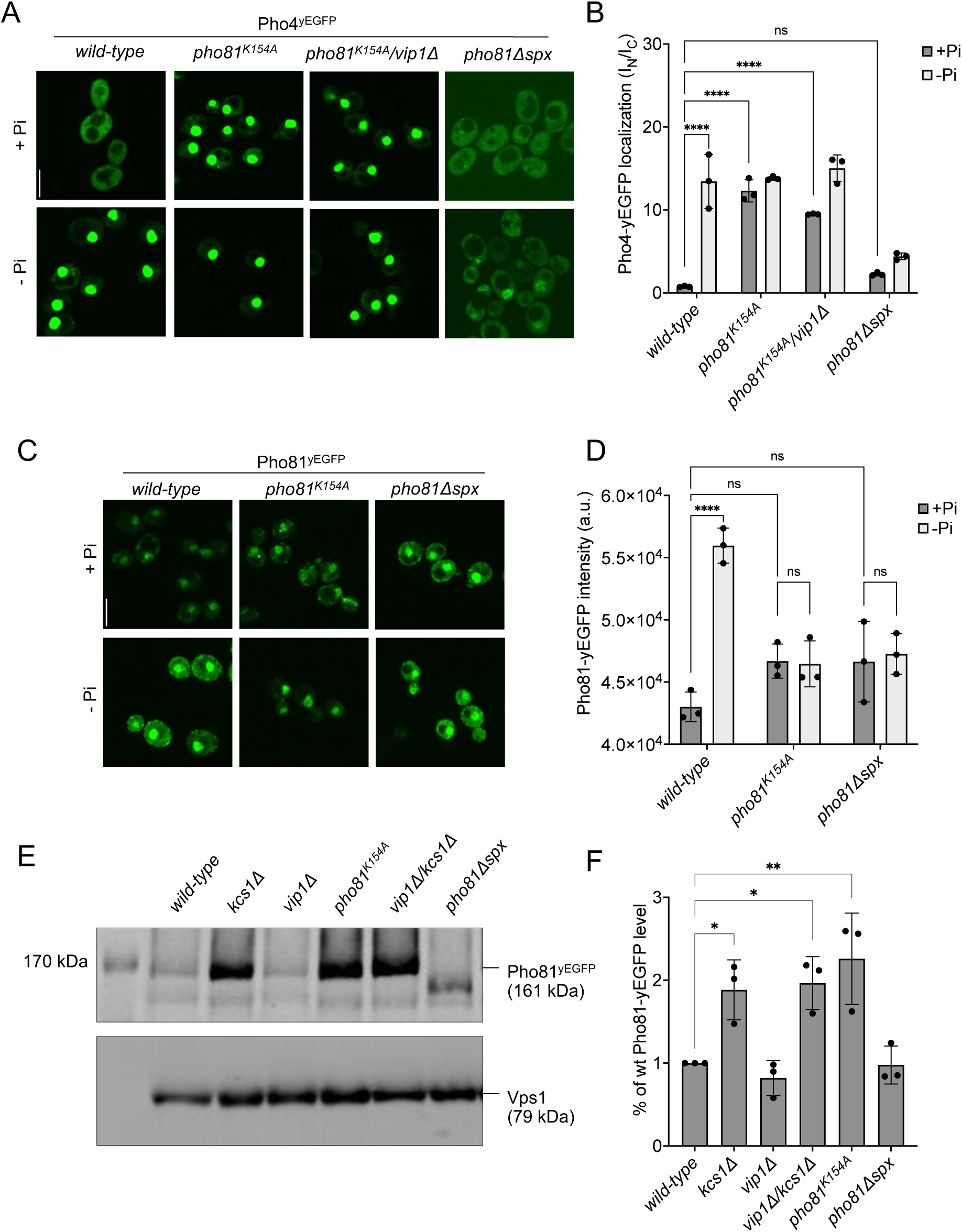
SPX-dependent activation of the PHO pathway. Cells were logarithmically grown in P_i_-replete SC medium as in Fig. 2A, washed, and incubated for further 4 h in medium with 7.5 mM of P_i_ (+ P_i_), or in starvation medium (- P_i_). (A) Pho4 relocation. The indicated cells expressed Pho4^yEGFP^ from its genomic locus. At the end of the 4 h starvation period, GFP fluorescence was imaged on a spinning disc microscope. Scale bar: 5 µm. λ_ex_: 488 nm; λ_em_: 500-530 nm. (B) Quantification of the nuclear localization of Pho4^yEGFP^ in images from A. Average intensity of Pho4^yEGFP^ fluorescence was determined by automated image segmentation and analysis. Pho4^yEGFP^ localization is quantified by the ratio of the average fluorescence intensities in the nucleus over the average fluorescence intensity in the cytosol (I_N_/I_C_). 100 to 200 cells were analyzed per condition and experiment. n=3. Means and standard deviation are indicated. (C) Pho81^yEGFP^ localization. The cells expressed the indicated variants of Pho81^yEGFP^ from its genomic locus. At the end of the 4 h growth period, GFP fluorescence was imaged as in A. (D) Quantification of the total cellular fluorescence of Pho81^yEGFP^. Images from C were subjected to automated segmentation and the average fluorescence intensity of the entire cells was quantified as in Fig. 3B. 100-200 cells were quantified per sample. n=3 experiments. Means and standard deviation are indicated. (E) Pho81^yEGFP^ expression assayed by Western blotting. Whole-cell protein extracts were prepared from cells expressing the indicated variants of Pho81^yEGFP^, which had been grown in P_i_-replete SC medium as in Fig. 2A. Proteins were analyzed by SDS-PAGE and Western blotting using antibodies to GFP. Vps1 was decorated as a loading control. (F) Quantification of Pho81^yEGFP^ blotting. Bands from experiments as in E were quantified on a LICOR Odyssey infrared fluorescence imager. The signals from wildtype cells were set to 1. Means and standard deviations are shown. n=3. For B, D, and F: **** p<0.0001; *** p<0.001; ** p<0.01; * p<0.05. n.s. not significant, determined with Turkey’s test.

Several lysines located on α-helix 4 of SPX domains form part of an inositol pyrophosphate binding patch (7, 12). Substituting them by alanine severely reduces the affinity of the domains for inositol pyrophosphates and can thus mimic the inositol pyrophosphate-free state. We determined the corresponding lysines in the SPX domain of Pho81 (Fig. S6), created a point mutation in the genomic PHO81 locus that substitutes one of them, K154, by alanine, and investigated the impact on the PHO pathway. Pho81^K154A-yEGFP^ should be correctly folded because it concentrates in the nuclei of the cells (49). *pho81^K154A^* cells constitutively activated the PHO pathway because Pho4^yEGFP^ accumulated in their nucleus already under P_i_-replete conditions (Fig. 5A, B). This constitutive activation was also observed in a *pho81^K154A^ vip1Δ* double mutant, confirming that 1-IP_7_ synthesis by Vip1 is not necessary for PHO pathway induction. Induction of the PHO pathway through SPX substitutions is further illustrated by *PHO81* expression, which itself is under the control of the PHO pathway (Fig. 5E, F). Western blots of protein extracts from cells grown under P_i_-replete conditions showed increased levels of Pho81 in *pho81^K154A^* cells, like those observed in strains that constitutively activate the PHO pathway due to the absence of IPPs, such as *kcs1Δ* and *vip1Δ/kcs1Δ*. They were significantly higher than in cells that do not constitutively activate the PHO pathway, such as wildtype or *vip1Δ*. In sum, these results suggest that the PHO pathway is regulated through binding of IPPs to the SPX domain of Pho81.

This critical role of the SPX domain for controlling Pho81 also consistent with the results from two random mutagenesis approaches (44, 52), which had received little attention in the previous model of PHO pathway regulation (19, 20). These random mutagenesis approaches generated multiple pho81^c^ alleles, which constitutively activate the PHO pathway and carry the substitutions G4D, G147R, 158-SGSG-159, or E79K. All these alleles affect the SPX domain at residues in the immediate vicinity of the putative IPP binding site (Figs. 6; S6). Three of the alleles (G4D, G147R, 158-SGSG-159) replace small glycine residues by bulky residues or insert additional amino acids, respectively, at sites where they should sterically interfere with binding of inositol pyrophosphates. We hence expect them to mimic the IPP-free state and constitutively activate the PHO pathway for this reason.

**Fig. 6:**
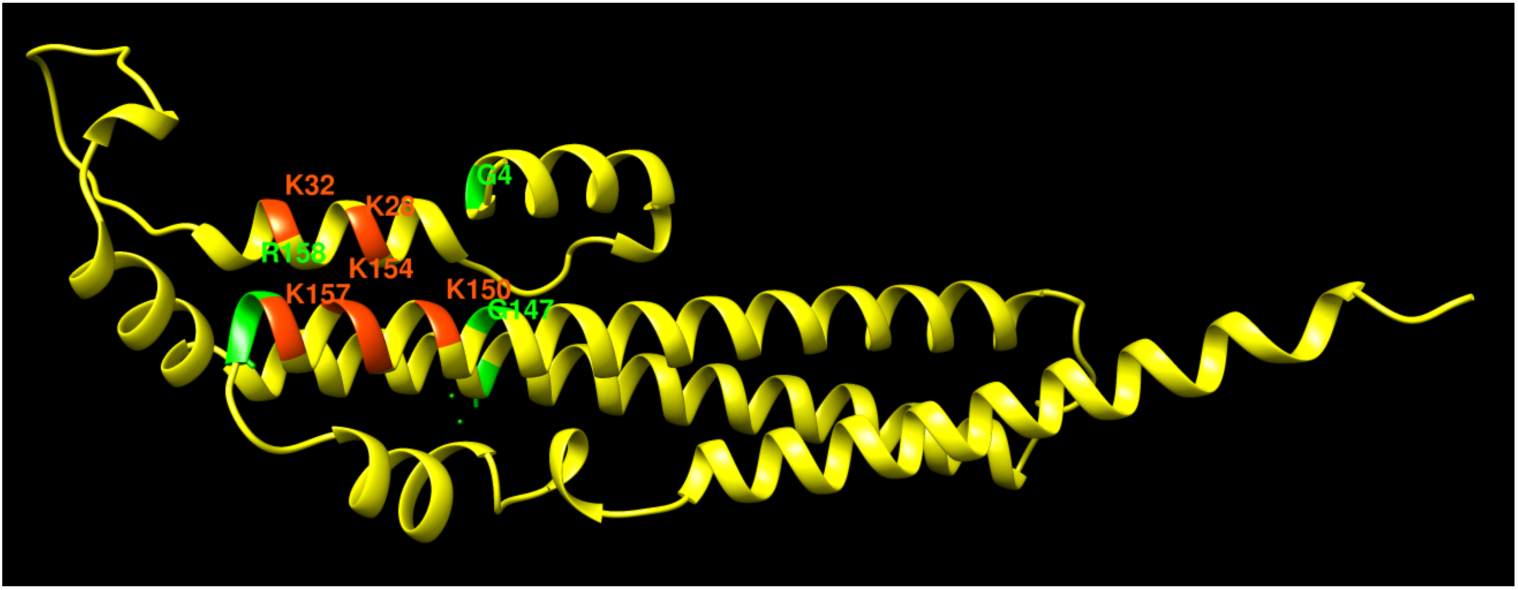
Pho81 residues leading to constitutive activation of the PHO pathway. The image shows an Alphafold prediction of the Pho81 SPX domain (amino acids 1-215), taken from Alphafold database model AF-P17442-F1 (81), in yellow. Basic residues of the putative IPP binding site have been identified by structure matching with the IP_6_-associated SPX domain of *VTC4* from *Chaetomium thermophilum* (PDB 5IJP). They are labeled in red. Residues from random mutagenesis screens, which lead to constitutive activation of the PHO pathway, are labeled in green.

### SPX and minimum domains contribute to the interaction of Pho81 with Pho85-Pho80 in vivo

The traditional model of PHO pathway activation has tied the effect of 1-IP_7_ to the minimum domain of Pho81, proposing that a complex of the two competitively inhibits Pho85 kinase (20). Since the results presented above argued against a role of 1-IP_7_ in inhibiting Pho85-Pho80 and triggering the PHO pathway, we explored potential alternative roles of the minimum domain by modeling the Pho80-Pho81 interaction with a Google Colab notebook for Alphafold multimer v3 (82). The non-templated structure prediction of the complex (Fig. 7) yields a Pho80 structure that agrees with an available crystal structure (83), showing R121 and E154 of Pho80 forming a salt bridge in a groove, which is critical for controlling Pho85 kinase (49). The prediction shows the groove binding a long unstructured loop of Pho81, which corresponds to the minimum domain. Substitutions in the minimum domain, and specifically of the residues binding the Pho80 groove in the prediction (residues 690-701), constitutively repress the PHO pathway when introduced into full-length Pho81, and they destabilize the interaction of Pho81 with Pho85-Pho80 (44, 49). The same effects are also shown by the *pho80^R121K^* and *pho80^E154V^* alleles, which ablate the critical salt bridge in the Pho80 groove (49, 83). The similarity of the effects of these substitutions validates the predicted interaction of the minimum domain with the R121/E154-containing groove. It invites a re-interpretation of the role of the minimum domain and suggests that this domain may serve as an anchor for Pho80 on Pho81 rather than acting directly on the Pho85 kinase.

**Fig. 7:**
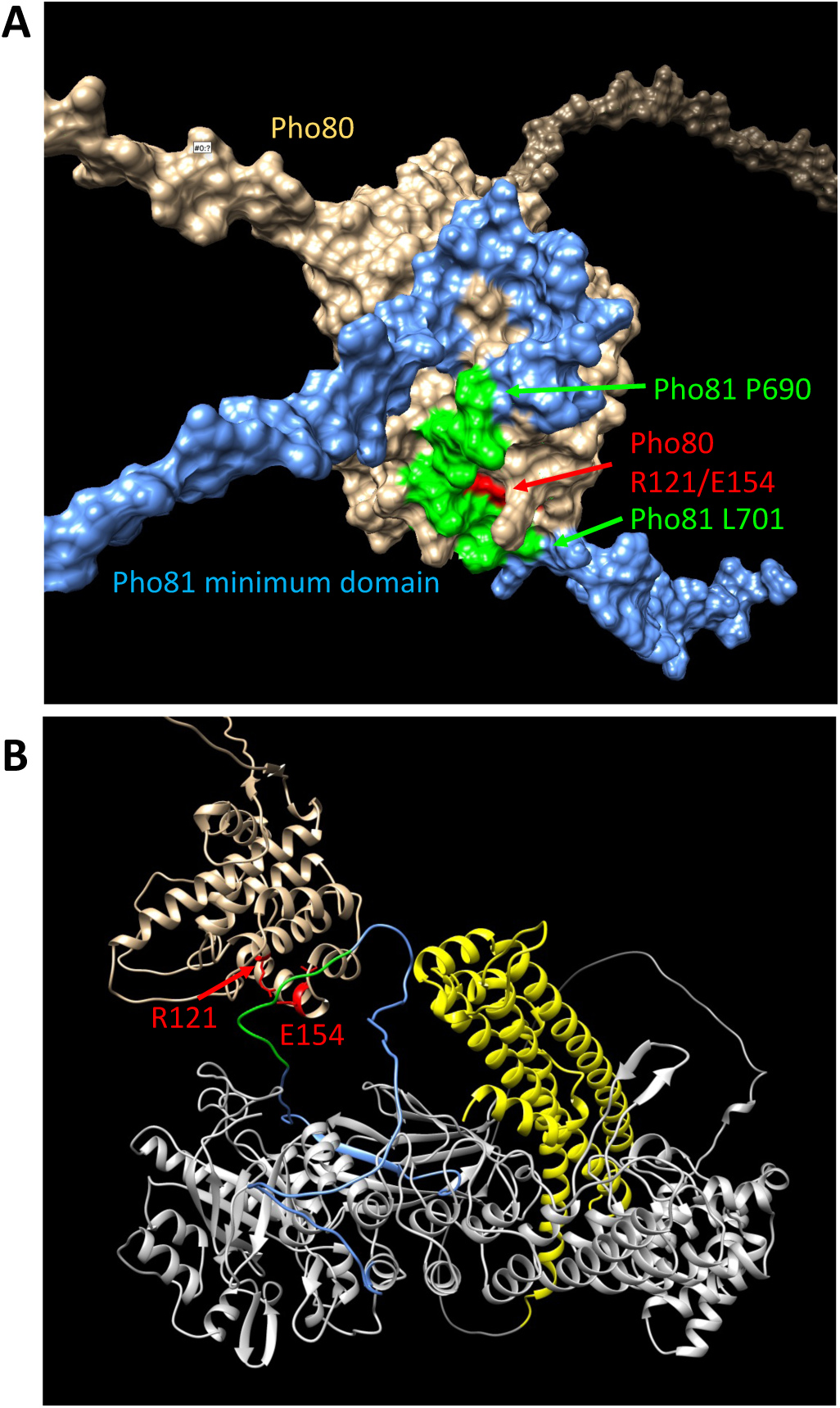
Structure predictions of the minimum domain in complex with Pho80. Alphafold multimer v3 was used to generate the following structure predictions: (A) Surface representation of Pho80 in complex with a peptide corresponding to the minimum domain of Pho81 (residues 645-724), showing the embedding of critical residues (690-701) of the minimum domain in a groove of Pho80 that contains the salt bridge between R121 and E154. (B) Ribbon representation of Pho80 in complex with full-length Pho81. Coloring: Beige - Pho80; red - E154, R121 of Pho80; blue: Pho81 minimum domain; green - central region of the Pho81 minimum domain that is critical for Pho85-Pho80 inhibition (aa 690-701); yellow: SPX domain of Pho81; grey - rest of Pho81 without assigned function. The complete dataset for both predictions has been deposited at Figshare under the DOI <u>10.6084/m9.figshare.c.6700281.</u>

The recruitment of Pho81 to the nucleus and its interaction with Pho85-Pho80 in wildtype cells is independent of the P_i_ regime, suggesting that it is dominated by a constitutive interaction (Fig. 5C) (49). This constitutive recruitment vanishes when the minimum domain binding site is compromised in *pho80^R121K^* and *pho80^E154V^* cells. Then, recruitment of Pho81 to the nucleus and to Pho85-Pho80 becomes P_i_-sensitive and is stimulated by P_i_ starvation (Fig. 8). We hence used *pho80^R121K^* and *pho80^E154V^* as tools to ask whether Pho81 might be tied to Pho85-Pho80 through a second layer of interaction that provides P_i_-dependence. As a *bona fide* P_i_-responsive element, the SPX domain of Pho81 was a prime candidate. We combined the *pho80^R121K^* and *pho80^E154V^* with substitutions in the putative IPP binding site of a fluorescent Pho81^yEGFP^ fusion protein, and with deletions of VIP1 or KCS1. Under P_i_-replete conditions, *pho80^R121K^* or *pho80^E154V^* cells showed Pho81^yEGFP^ in the cytosol, but they accumulated the protein in the nucleus only upon P_i_ starvation (Fig. 8). This rescue by starvation was more pronounced in *pho80^R121K^* than in *pho80^E154V^*. Pho81^K154A-yEGFP^, with a substitution in its putative IPP binding site designed to mimic the low-P_i_ state, accumulated in the nucleus of *pho80^R121K^* and *pho80^E154V^* cells already under P_i_-replete conditions, i.e., it behaved constitutively as under P_i_ starvation. *vip1Δ* mutants behaved like the wildtype, accumulating Pho81^yEGFP^ in the nucleus *of pho80^R121K^* and *pho80^E154V^* cells upon P_i_ starvation, whereas *kcs1Δ* cells showed nuclear Pho81^yEGFP^ already under P_i_-replete conditions. Thus, interfering with IPP signaling, be it through P_i_ starvation, ablation of IPP synthesis, or substitution of the putative IPP binding site of Pho81, partially compensated for the destabilization of the minimum domain binding site in Pho80.

**Fig. 8.**
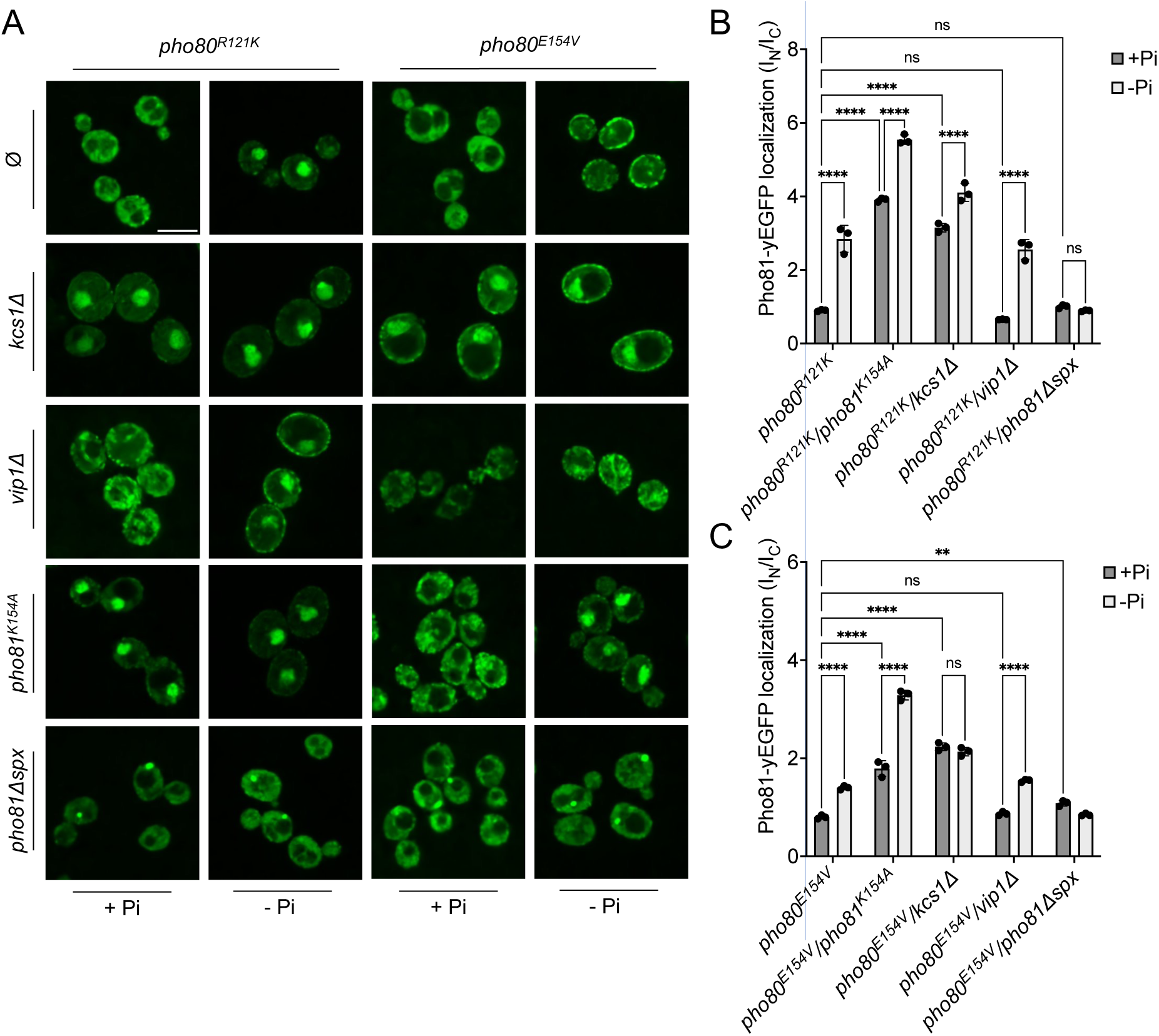
**Localization of Pho81^yEGFP^ in *pho80^R121K^* and *pho80^E154V^* loss-of-affinity mutants**. Cells were logarithmically grown in P_i_-replete SC medium as in Fig. 2A, washed, and incubated for further 4 h in medium with 7.5 mM of P_i_ (+ P_i_), or in starvation medium (- P_i_). (A) Pho81 imaging. Cells expressing the indicated variants of Pho81^yEGFP^ and Pho80 from their genomic loci were imaged on a spinning disc microscope after 4 h of growth in the presence of 7.5 mM Pi (+ P_i_), or after 4 h of P_i_ starvation (- P_i_). Scale bar = 5 μM. λ_ex_: 488 nm; λ_em_: 500-530 nm. Note that the fluorescent dots visible in *pho81^Δspx^*cells are not nuclei because they are too small and at positions where nuclei are not found. Their location was not further investigated because it is not essential for this study. (B) Quantification of the nuclear localization of Pho81^yEGFP^in *pho80^R121K^* cells. Images from A were subjected to automated segmentation and quantification of the average fluorescence intensity in the nucleus and cytosol as in Fig. 3B. 100-200 cells were quantified per sample. n=3 experiments. (C) Quantification of the nuclear localization of Pho81^yEGFP^ in *pho80^E154V^* mutants was performed as in B. For B and C: **** p<0.0001; *** p<0.001; ** p<0.01; * p<0.05; n.s. not significant, determined with Turkey’s test.

Two *in vivo* observations are consistent with the notion that this compensation may reflect an interaction of the SPX domain with Pho85-Pho80: First, Pho81^ΔSPX-yEGFP^, which lacks this SPX domain, did not show starvation-induced nuclear accumulation in *pho80^R121K^* and *pho80^E154V^* cells (Fig. 8). Second, a yEGFP fusion of only the SPX domain of Pho81 (Pho81SPX^yEGFP^), which lacks the rest of the protein and thereby excludes a constitutive interaction through the minimum domain, localized mainly to the cytosol of PHO80 wildtype cells under P_i_-replete conditions, but it concentrated in the nucleus upon P_i_ starvation (Fig. S7). This concentration was not observed in *pho80Δ* cells, which lack nuclear Pho85-Pho80. By contrast, *pho80^R121K^* or *pho80E^154V^* cells, which only target the minimum domain-Pho80 interaction but leave Pho85-Pho80 kinase active and in the nucleus (83), retained the capacity to accumulate Pho81SPX^yEGFP^ in their nuclei upon P_i_ starvation. These results suggest that the SPX-domain contributes to the interaction of Pho81 and Pho85-Pho80. Our observations are consistent with a model in which this SPX-Pho85-Pho80 interaction is independent of the minimum domain but controlled by P_i_ through IPPs, which can destabilize it and relieve the inhibition of Pho85-Pho80.

## Discussion

The transcriptional response of *S. cerevisiae* to varying P_i_ supply is an important model system for studying gene regulation, promotor structure, and the impact of chromatin structure (42, 84). However, identification of the inositol pyrophosphates relaying phosphate availability to this PHO pathway has been challenging. This is mainly due to limitations in analytics. The recent development of a CE-MS method has provided a potent tool, allowing IPP analysis without radiolabeling and under any physiological condition. The improved separation, higher throughput and lower cost of this approach led us to re-investigate the role of IPPs in PHO pathway regulation. The results inspire a revised model that differs from the traditional hypothesis in three aspects: We propose that P_i_ scarcity is signaled by a decrease in IPPs rather than by an increase; a critical function is ascribed to the loss of 1,5-IP_8_ instead of 1-IP_7_; and we postulate that the N-terminal SPX domain of Pho81 rather than the central minimum domain is the primary receptor mediating IPP-control over the PHO pathway.

Our observations reveal an effect that misled the interpretation of several preceding studies. This includes our own work on the activation of the polyphosphate polymerase VTC (85), which is controlled by SPX domains (4, 6, 12, 86). Our *in vitro* activity assays had shown a clear order of potency of IPPs in stimulating VTC, with 1,5-IP_8_ showing an apparent EC_50_ 30-fold below that of 5-IP_7_ and 50-fold below that of 1-IP_7_ (4). In this work, we nevertheless dismissed 1,5-IP_8_ as a physiological stimulator of the VTC SPX domains, based on the argument that a *vip1Δ* mutant, which lacks the enzyme for 1-IP_7_ and 1,5-IP_8_ synthesis, shows quite normal polyP accumulation in vivo. We proposed 5-IP_7_ for this role because a *kcs1Δ* mutant, which lacks the IP6K making this compound, also lacks polyP. The IPP analyses that we present now call for a re-interpretation of this data. The presence of polyP in a *vip1Δ* mutant can well be explained by the 20-fold over-accumulation of 5-IP_7_ in this mutant, which can compensate for the lower EC_50_ and permit stimulation of VTC through 5-IP_7_. In wildtype cells, however, where 5-IP_7_ is only twice as abundant as 1,5-IP_8_ (Fig. 2), 1,5-IP_8_ should be the relevant activator of VTC for polyP synthesis under P_i_-replete conditions.

Then, the situation is also compatible with results from *S. pombe*, where ablation of the Vip1 homolog Asp1 impairs polyP synthesis (24, 87). The transcriptional phosphate starvation response in *S. pombe* also depends on IPPs. However, the downstream executers differ from those present in *S. cerevisiae*. Fission yeast contains no homologs of Pho81, and the responsible transcription factor is not controlled through a Pho85-Pho80-like kinase (32–34). P_i_-dependent transcription strongly depends on lncRNA-mediated interference, which may be linked to IPPs through an SPX-domain containing ubiquitin E3 ligase (30, 35, 36, 87). Nevertheless, important basic properties of the IP6K and PPIP5K system are conserved. Like in baker’s yeast, deletion of the Vip1 homolog Asp1 leads to a strong increase in IP_7_ (27). Characterization of the purified enzyme also showed that P_i_ stimulates net production of IP_8_ by inhibiting the phosphatase domain of Asp1 (31). In line with this, we observed that IP_8_ rapidly decreased upon P_i_ starvation also in *S. pombe* (Fig. S3). As for Vip1 in baker’s yeast, genetic ablation of Asp1 does not interfere with the induction of the phosphate-regulated gene *PHO1* by P_i_ starvation, suggesting that 1-IP_7_ and 1,5-IP_8_ are not essential for this transcriptional response (32). In P_i_-replete media, however, ablation of Asp1 hyper-represses *PHO1*, and putative overproduction of 1-IP_7_ and/or 1,5-IP_8_ by selective inactivation of only the Asp1 phosphatase domain induces this gene (32, 88). It remains to be seen whether these discrepancies originate from a similar disequlibration of IP_7_ species as in *S. cerevisiae* (ablation of Asp1 increases the IP_7_ pool 4- to 5-fold (27)), or whether they are due to other processes such as lncRNA-mediated interference, which plays an important role in P_i_ homeostasis in *S. pombe* (36–38).

The PHO pathway in *Cryptococcus neoformans*, a pathogenic yeast, shares many protein components with that of *S. cerevisiae* and might hence be expected to function in a similar manner (89). Yet a recent study on this organism proposed 5-IP_7_ as the critical inducer of the PHO pathway (21). This model relies on the observation that *kcs1Δ* mutants, which lack 5-IP_7_, cannot induce the PHO pathway in this yeast. This is the opposite behavior compared to *S. cerevisiae*. It was hence concluded that, despite an experimentally observed moderate decline in 5-IP_7_ upon short P_i_ starvation, the remaining pool of 5-IP_7_ was required for PHO pathway activity. Substitution of basic residues in the IPP-binding pocket of *C. neoformans* Pho81 interfered with PHO pathway activation, lending further support to the conclusion. However, the same study also showed that Pho81 was lacking completely from kcs1Δ mutants. When the Pho81 IPP binding site was compromised by amino acid substitutions the Pho81 protein level was significantly reduced, and the protein almost vanished upon P_i_ starvation. This offers an alternative explanation of the effects because absence of the critical inhibitor Pho81 activates Pho85/80 kinase and represses the PHO pathway independently of P_i_ availability (43, 44). Destabilization of Pho81 will then mask effects of IPPs. Thus, it remains possible that the PHO pathway in *C. neoformans* is controlled as in *S. cerevisiae*, i.e. that P_i_ starvation is signaled by a decline of 1,5-IP_8_. In line with this, we observed that *C. neoformans* and *S. pombe* cells lose all IPPs within 2 hours of phosphate starvation (Fig. S3). As in *S. cerevisiae*, 1,5-IP_8_ showed a faster and more profound response. Therefore, we consider it as likely that the loss of 1,5-IP_8_ signals P_i_ starvation also in *C. neoformans* and *S. pombe*.

Our results revise the widely accepted model of increased 1-IP_7_ as the trigger of the PHO pathway and of the minimum domain of Pho81 as its receptor (19, 20). This model rests upon an increase of 1-IP_7_ upon P_i_ starvation, a reported constitutive activation of the PHO pathway in a *ddp1Δ* mutant, and upon the lack of rapid PHO pathway induction in a *vip1Δ* mutant. A later study from O’Shea and colleagues revealed that *vip1Δ* cells finally induced the PHO pathway after prolonged P_i_ starvation (90), but this did not lead to a revision of the model. Our results rationalize the strong delay of *vip1Δ* in PHO pathway induction by the overaccumulation of 5-IP_7_ in this mutant, which leads to aberrant PHO pathway silencing through this compound. Under P_i_ starvation, it takes several hours longer to degrade this exaggerated pool, which provides a satisfying explanation for the delayed PHO pathway induction in *vip1Δ* cells. Even though earlier studies, due to the limited resolution of the radio-HPLC approach, could not differentiate between 1- and 5-IP_7_, these studies reported an increase in the IP_7_ pool upon P_i_ starvation (19). Other studies using a similar radio-HPLC approach suggested a decline of the IP_7_ pool upon long P_i_ withdrawal, but they analyzed only a single timepoint and could not exclude an earlier increase in 1-IP_7_ as a trigger for the PHO pathway (6, 21, 22). In our CE-MS analyses, we consistently observed strong, continuous declines of 1,5-IP_8_, 5-IP_7_ and 1-IP_7_ and, upon several hours of P_i_ starvation, their almost complete disappearance. Furthermore, we could not confirm the reported constitutive activation of the PHO pathway in a *ddp1Δ* mutant (19), although this mutant did show the expected increased 1-IP7 concentration. These observations argue against a role of 1-IP_7_, 5-IP_7_ or 1,5-IP_8_ as activators for Pho81 and the PHO pathway.

The link between 1-IP_7_ and Pho81 activation relied mainly on the use of the 80 amino acid minimum domain from the central region (19, 20, 49). This domain was used in *in vitro* experiments, where 1-IP_7_ stimulated binding of the minimum domain to Pho85-Pho80 and inhibition of the enzyme. Given that 1-IP_7_ disappears under P_i_ starvation it appears unlikely that this *in vitro* effect is physiologically relevant. Furthermore, the apparent dissociation constant of the complex of 1-IP_7_ with the minimum domain and Pho80-Pho85 was determined to be 20 µM (20). The minimum domain is thus unlikely to respond to 1-IP_7_ in the cells because its K_d_ is almost 50-fold above the cellular concentrations of 1-IP_7_ in P_i_-replete medium (0.5 µM; Fig. 2), and at least 3 orders of magnitude above the concentration remaining on P_i_ starvation media (< 20 nM). Another discrepancy is that the *in vitro* inhibition of Pho85-Pho80 by the minimum domain was reported to be specific for 1-IP_7_ whereas 5-IP_7_ had no effect (20). This contrasts with the *in vivo* situation, where we find that even massive overaccumulation of 1-IP_7_ by ablation of Ddp1 cannot trigger the PHO pathway, but where suppression of 5-IP_7_ and 1,5-IP_8_ production by ablation of Kcs1 does activate it constitutively (Fig. 3)(23).

An argument supporting a role of the minimum domain in mediating P_i_-responsiveness was provided by *in vivo* experiments in a *pho81Δ* mutant, where P_i_-dependent activation of the PHO pathway could be partially rescued through over-expression of the minimum domain (44, 49). PHO pathway activation was assayed through *PHO5* expression. This is relevant because the *PHO5* promotor is not only controlled through Pho85-Pho80-mediated phosphorylation of Pho4, but also through changes in nucleosome occupancy, which substantially influence activity and activation threshold of the *PHO5* promotor (77, 79). It is thus conceivable that constitutive overexpression of the minimum domain reduces Pho85-Pho80 activity moderately, but sufficiently to allow regulation of *PHO5* expression through nucleosome positioning, which changes in a P_i_-dependent manner (77).

Even though the minimum domain is unlikely to function as a receptor for IPPs this does not call into question that the minimum domain is critical for Pho81 function. A role of this domain is supported by substitutions and deletions in this domain, introduced into full-length Pho81, which constitutively repress the PHO pathway (44, 49). Our modeling approach provides a satisfying link of these residues to Pho85-Pho80 regulation. It suggests the minimum domain to be an extended unstructured loop that binds Pho80 at a groove that encompasses R121 and E154, residues that emerged from a genetic screen as essential for recruiting Pho81 to Pho85-Pho80 and for inhibiting the kinase (49). Therefore, we propose that the minimum domain does provide a critical connection between Pho81 and Pho80, which is per se IPP-independent, whereas the SPX domain of Pho81 may create additional contacts to Pho80-Pho85 and impart regulation through IPPs.

Taking all these aspects together, we propose the following working model (Fig. 9): P_i_ sufficiency in the cytosol is signaled through the accumulation of 1,5-IP_8_. 1,5-IP_8_ binding to the SPX domain of Pho81 labilizes the interaction of this domain with Pho85-Pho80, allowing the kinase to become active. The now un-inhibited kinase phosphorylates Pho4, triggers its relocation into the cytosol and represses the PHO pathway. P_i_ limitation is signaled by a decline in 1,5-IP_8_, resulting in the ligand-free form of the SPX domain and the activation of Pho81. Inhibition of Pho85-Pho80 requires the Pho81 minimum domain to bind a groove in the cyclin Pho80, stabilizing the complex and inhibiting Pho85 kinase. An obvious question arising from this model is whether the Pho81 SPX domain itself could activate Pho85-Pho80 by directly binding these proteins. Alternatively, or in addition, Pho85-Pho80 might be controlled through the availability of the minimum domain for Pho80 binding, which might be restricted through an IPP-dependent SPX-minimum domain interaction. These aspects will be explored in future work.

**Fig. 9:**
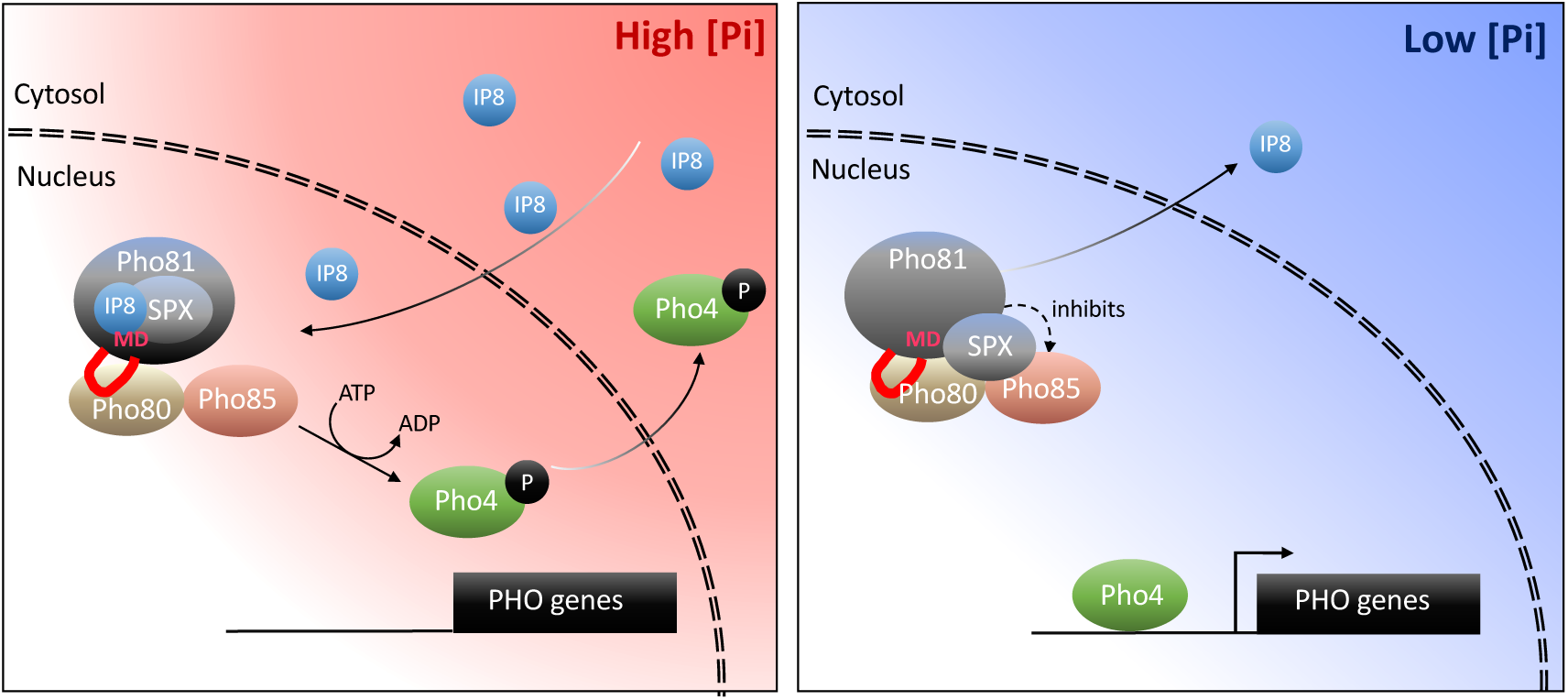
Working model on the control of the PHO pathway through 1,5-IP_8_ and Pho81. At high P_i_ concentrations, inositol pyrophosphates accumulate. 1,5-IP_8_ binds the SPX domain of Pho81, which labilizes the interaction of Pho81 with Pho80 and prevents Pho81 from inhibiting Pho85-Pho80 kinase. In low P_i_, 1,5-IP_8_ declines. Liberation of the SPX domain from this ligand allows this domain to interact with Pho85-Pho80. This, and the interaction of the minimum domain with a critical groove of Pho80, allow Pho81 to inhibit Pho80-85. The resulting dephosphorylation of Pho4 triggers its concentration in the nucleus and activation of the PHO pathway. This inhibition is reinforced by increased expression of PHO81, which itself is a PHO pathway-controlled gene.

This revision of PHO pathway regulation permits to develop a coherent picture on intracellular phosphate signaling across the eukaryotic kingdoms. Phosphate abundance being indicated by the accumulation of 1,5-IP_8_ is compatible with the situation in mammalian cells, where 1,5-IP_8_ triggers P_i_ export through the putative phosphate transporter XPR1 (14, 91), and with the situation in plants, where IP_8_ is particularly abundant upon shift to high-P_i_ substrate and inhibits transcription factors for the phosphate starvation response (7, 10, 16, 18). Thus, loss of 1,5-IP_8_ may generally signal phosphate scarcity in eukaryotic cells.

## Materials and Methods

### Yeast culture

Unless stated otherwise, all experiments were performed with *Saccharomyces cerevisiae* BY4741 cells or derivatives thereof. Cells have been grown in liquid Synthetic Complete (SC) medium of the following composition: 3.25 g yeast nitrogen base (YNB) without amino acids (Difco 291920); 1 g Synthetic Complete Mixture (Kaiser) Drop-Out: Complete (Formedium DSCK1000); 50 ml 20% glucose; 450 ml demineralized water. Phosphate starvation experiments were performed in SC-P_i_ medium: 3.25g YNB without amino acids and phosphate, supplemented with KCl (Formedium CYN6703); 1 g Synthetic Complete Mixture (Kaiser) Drop-Out: Complete (Formedium DSCK1000); 50ml 20% glucose; 450 ml water. These synthetic media were sterile filtered and tested for absence of free P_i_ by the malachite green assay. Cultures were shaken in capped Erlenmeyer flasks at 150 rpm at 30°C unless stated otherwise.

### Yeast strains and their genetic manipulation

The *S. cerevisiae* strains used in this study are listed and described in Supplementary Information. DNA transformations were carried out with overnight cultures at 30°C in YPD medium (Yeast Extract-Peptone-Dextrose) using the following protocol: Cultures (not exceeding 10^7^ cells/mL) were centrifuged at 1800 × g for 2 min. 25 μL of pellet was resuspended in 25 μL of LiAc-TE 0.1 M (Lithium acetate 0.1M, Tris 10 mM, EDTA 1 mM). The cell suspension was supplemented with 240 μL of PEG 50%, 36 μL of LiAC 1 M, 5 μL of boiled ssDNA, and the DNA (200 ng of plasmid and/or 0.5-2 mg of PCR product). After 30 min at 30°C, a heat shock was carried out at 42°C for 20 min. Cells were then centrifuged at 1800 × g for 2 min, resuspended in 150 μL of water, and grown on selective plates for 2 to 4 days. Fluorescent protein tags have been obtained using a standard method (92), except for Hta2 tagging, which was performed by CRISPR-Cas9. All introduced single mutations, knockouts, and fluorescent protein tags were confirmed by PCR and sequencing.

### Plasmid constructions

The plasmids used to carry out CRISPR-Cas9 genome editing were obtained by cloning hybridized oligonucleotides coding for the sgRNA between the XbaI and AatII restriction sites of the parent plasmid (pSP473, pSP475 or pSP476). The plasmids used to overexpress *VIP1* (pVC97 and pVC114) were obtained by cloning the *VIP1* open reading frame between the BamHI and SalI restriction sites of the parent plasmid (respectively pRS415-GPD and pRS413-GPD) (93). The plasmid used to monitor *prPHO5-GFP* expression (pVC115) was obtained by replacing the *pgk1* promoter with the -1 to -1500 region of the *PHO5* promoter and by replacing the gene for G418 resistance to the coding region of the *LEU2* gene within the pCEV-G1-Km backbone (Addgene).

### Fluorescence microscopy

Cells were grown logarithmically in SC medium for 12-15 h until they reached a density of 10^7^ cells/mL. For P_i_ starvation experiments, overnight-grown cells were washed twice, resuspended in SC-P_i_ medium at a density of ca. 5×10^6^ cells/mL, and incubated at 30°C. Fluorescence images were recorded on a Nikon Eclipse Ti2/Yokogawa CSU-X1 Spinning Disk microscope equipped with two Prime BSI sCMOS cameras (Teledyne Photometrics), a LightHUB Ultra® Laser Light (Omicron Laserage), and an Apo TIRF 100x/1.49 Oil lens (Nikon). Time-lapse experiments were performed on a Nikon Eclipse Ti2 inverted microscope equipped with a Prime BSI sCMOS camera (Teledyne Photometrics), a Spectra X Light Engine (Lumencor), a Plan Apo λ 100x/1.45 Oil lens (Nikon). Microscopy chambers were made using sticky-Slides VI ^0.4^ (Ibidi) glued onto concanavalin a-coated cover slips (Knittel glass, 24 × 60 × 0.13 – 0.17 mm). The cell suspension (200 μL, 10^7^ cells/mL) was added to the chamber. After 30 seconds of sedimentation, a flow of SC medium was applied using the microfluidic flow controller AF1 Mk2 (Elveflow) coupled to a MUX distributor (Elveflow). After 2 min, the flow was switched to SC-P_i_ medium, and imaging was started. After further 2 min, the flow was stopped, and the time lapse was continued by recording one image every 5 min. The temperature was kept at 30°C using the stage-top incubator Uno-Controller (Okolab).

### Western Blots

5 mL of cells grown overnight in SC medium (until ca. 10^7^ cells/mL) was centrifuged (1’800 × g, 2 min). Cells were resuspended in 1 mL of lithium acetate (2 M) and centrifuged as before. Cells were resuspended in 1 mL of sodium hydroxide (0.4 M) and kept on ice for 5 min. After centrifugation (1’800 × g, 2 min), cells were resuspended in 100 μL of SDS-PAGE Sample Buffer and boiled for 5 min. After centrifugation under the same conditions, the supernatant was collected, and the protein content was measured using a NanoDrop 1000 Spectrophotometer (Witec). Supernatant concentrations were adjusted, and the samples were loaded on SDS-polyacrylamide gels.

### Inositol pyrophosphate quantification

Cells were logarithmically grown in SC medium overnight to reach a density of 10^7^ cells/mL. 1 mL of cell culture was mixed with 100 µL of 11 M perchloric acid (Roth HN51.3). After snap-freezing in liquid nitrogen, the samples were thawed and cell debris was removed by centrifugation (15’000 × g, 3 min, 4°C). Titanium dioxide beads (Titansphere TiO_2_ beads 5 mm, GL Sciences 5020-75000) were pretreated by washing them once with water, once with 1 M PA and finally resuspending them in 200 µl 1 M perchloric acid. The cell supernatant was mixed with 1.5 mg of beads and incubated with shaking or on a rotating wheel for 15 min at 4°C. The beads were collected by centrifugation (15’000 × g, 1 min, 4°C) and washed twice with 1 M perchloric acid. Inositol phosphates were eluted by 300 µL of 3% ammonium hydroxide and the eluate was evaporated using a speed-vac concentrator at 42°C and 2000 rpm overnight. The pellet was resuspended using 20 µl of water. CE-ESI-MS analysis was performed on an Agilent 7100 CE coupled to triple quadrupole mass spectrometer (QqQ MS) Agilent 6495c, equipped with an Agilent Jet Stream (AJS) electrospray ionization (ESI) source. Stable CE-ESI-MS coupling was enabled by a commercial sheath liquid coaxial interface, with an isocratic liquid chromatography pump constantly delivering the sheath-liquid.

### Spectrofluorimetry

Cells were grown overnight to a density of 5×10^6^ cells/mL in SC medium. They were then resuspended in SC-P_i_ medium after two washing steps. For each timepoint, the concentration of the cell suspension was measured and adjusted to 10^7^ cells/mL before fluorescence measurement. The fluorescence of yEGFP (λ_ex_: 400 nm; λ_em_: 500-600 nm; λ_cutoff_: 495 nm) and mCherry (λ_ex_: 580 nm; λ_em_: 600-700 nm; λ_cutoff_: 590 nm) were measured on a Molecular Devices SpectraMax Gemini EM microplate fluorimeter at 30°C.

### Analysis of IPP in C. neoformans and S. pombe

*C. neoformans grubii* wildtype cells were grown in liquid synthetic complete (SC) medium to logarithmic phase. They were sedimented (3’000 × g, 1 min, 4°C), washed twice with phosphate-free medium, aliquoted, and used to inoculate four 25 ml cultures in SD medium lacking phosphate. The inoculum was adjusted (starting OD_600_ at 0.9, 0.6, 0.4, 0.3 etc, respectively) such that, after further incubation for 0 to 240 min, all samples had similar OD_600_ at the time of harvesting. The sample for the 0 min timepoint was taken from the culture in SC medium. The starvation time was counted beginning with the first wash. At the indicated time points, the OD_600_ of the culture was measured. A 4 ml sample was collected, supplemented with 400 µl of 11 M perchloric acid, frozen in liquid nitrogen and kept at - 80°C. The samples were thawed and IPPs were extracted using 6 mg TiO_2_ beads due to the higher number of cells. For *S. pombe*, wildtype cells were prepared in a similar way to *C. neoformans.* 3 ml of culture was collected, supplemented with 300 µl of 11 M perchloric acid, and frozen in liquid nitrogen. All further steps were as described above for *S. cerevisiae* samples.

## Acknowledgements

We thank Julianne Djordjevic, Sophie Martin and John York for strains and discussion, Dorothea Fiedler for ^13^C-labelled IPP standards, and Andrea Schmidt for technical assistance. This study was supported by grants from SNSF (CRSII5_170925) and ERC (788442) to AM, from HFSP (LT000588/2019) to G-DK, and from the German Research Foundation (DFG) under Germanýs Excellence Strategy (CIBSS – EXC-2189 – Project ID 390939984) to HJJ.

## Supplementary Information

### Strains

The *S. cerevisiae* strains used in this study were all obtained by genetic manipulations from the BY4741 background described in Table S1. This strain corresponds to the wild-type of this study. For the sake of clarity, the BY4741 background was therefore not indicated in the genotype of each mutant strain.

**Table S1.**
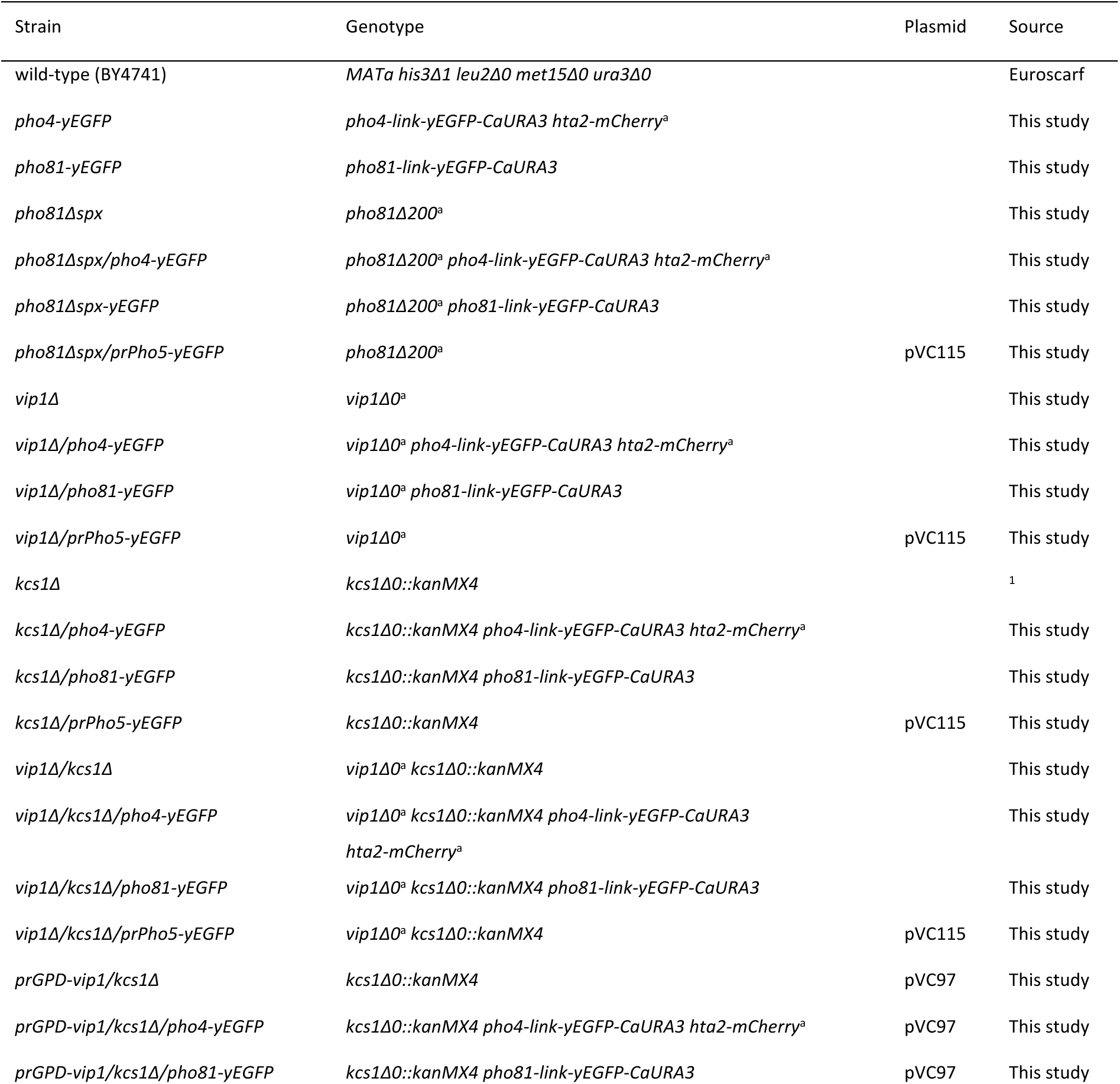
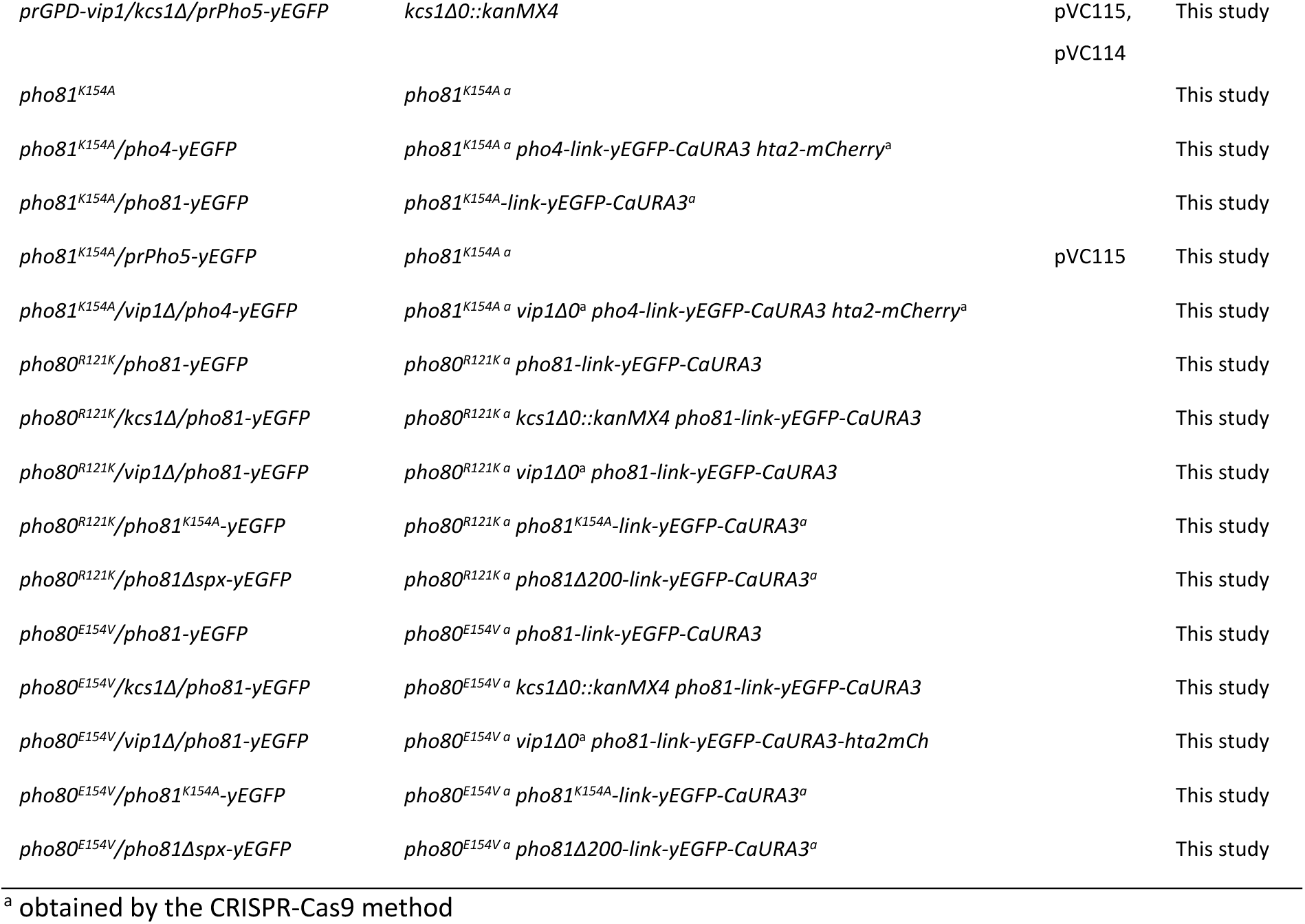
Strains used in this study

### Plasmids

**Table S2.** Plasmids used in this study

| Plasmid | Description | Source |
| --- | --- | --- |
| pVC115 | pCEV-G1-LEU- *prPHO5-yegfp prTEF-mcherry leu2* | This study |
| pVC97 | pRS415 *prGPD-vip1 leu2* | This study |
| pVC114 | pRS413 *prGPD-vip1 his3* | This study |
| pED109 | *pGTL-mcherry* | Serge Pelet |
| pSP473 | pRS423 *prGPD-cas9 his3* | Serge Pelet |
| pSP475 | pRS425 *prGPD-cas9 leu2* | Serge Pelet |
| pSP476 | pRS426 *prGPD-cas9 ura3* | Serge Pelet |
| pSP478 | pRS425 *prGPD-cas9_sgRNA_hta2 leu2* | Serge Pelet |
| pVC24 | pRS425 *prGPD-cas9_sgRNA_pho81 leu2* | This study |
| pVC43 | pRS425 *prGPD-cas9_sgRNA_kcs1 leu2* | This study |
| pVC44 | pRS423 *prGPD-cas9_sgRNA_vip1 his3* | This study |
| pVC110 | pRS425 *prGPD-cas9_sgRNA_vip1 leu2* | This study |
| pVC72 | pRS423 *prGPD-cas9_sgRNA_pho80_1 his3* | This study |
| pVC73 | pRS423 *prGPD-cas9_sgRNA_pho80_2 his3* | This study |
| pVC108 | pRS425 *prGPD-cas9_sgRNA_pho80_1 leu2* | This study |
| pVC109 | pRS425 *prGPD-cas9_sgRNA_pho80_2 leu2* | This study |
| pVC119 | pRS315 *hta2-mCherry* | This study |

### Oligonucleotides

**Table S3.**
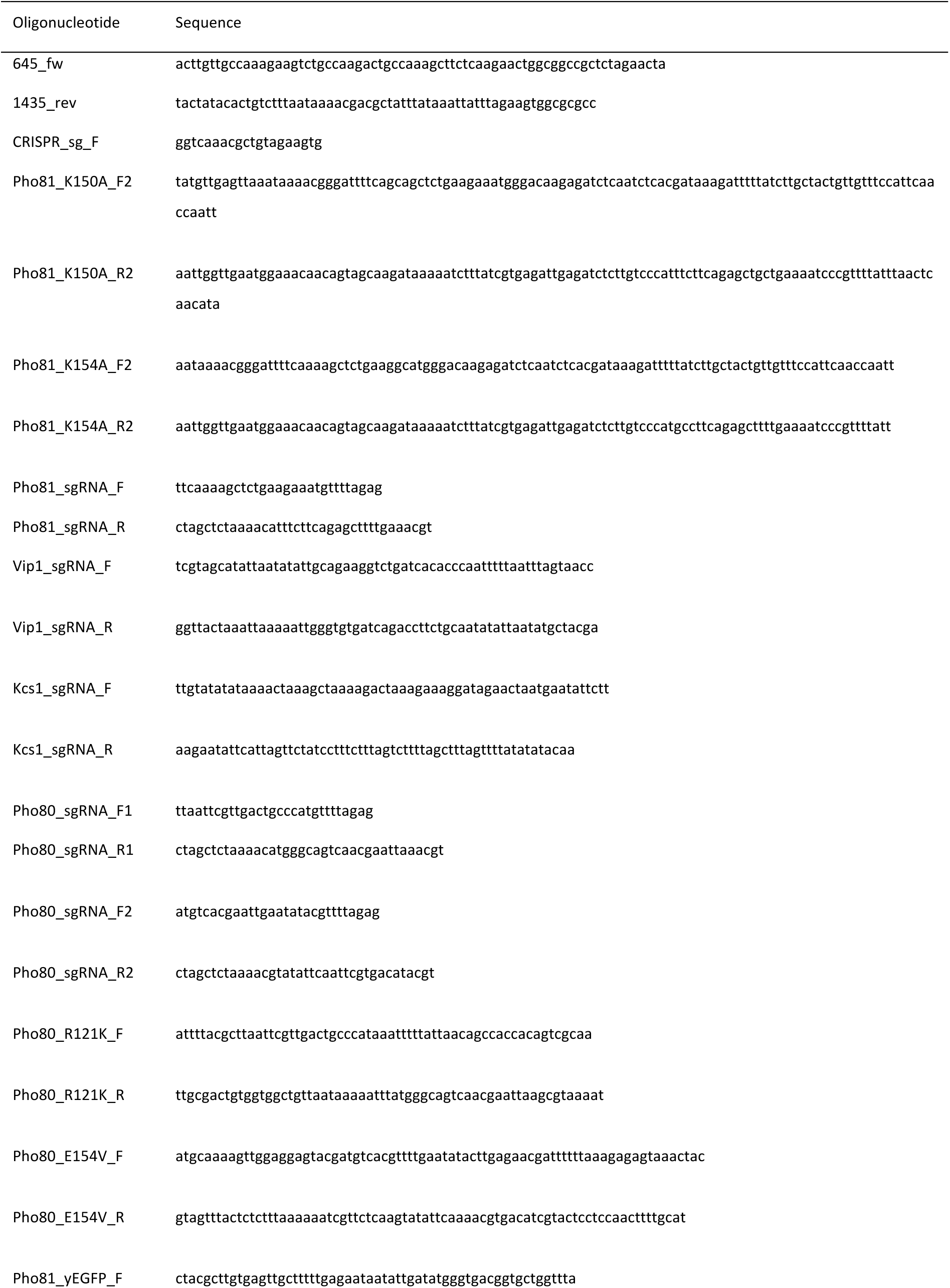
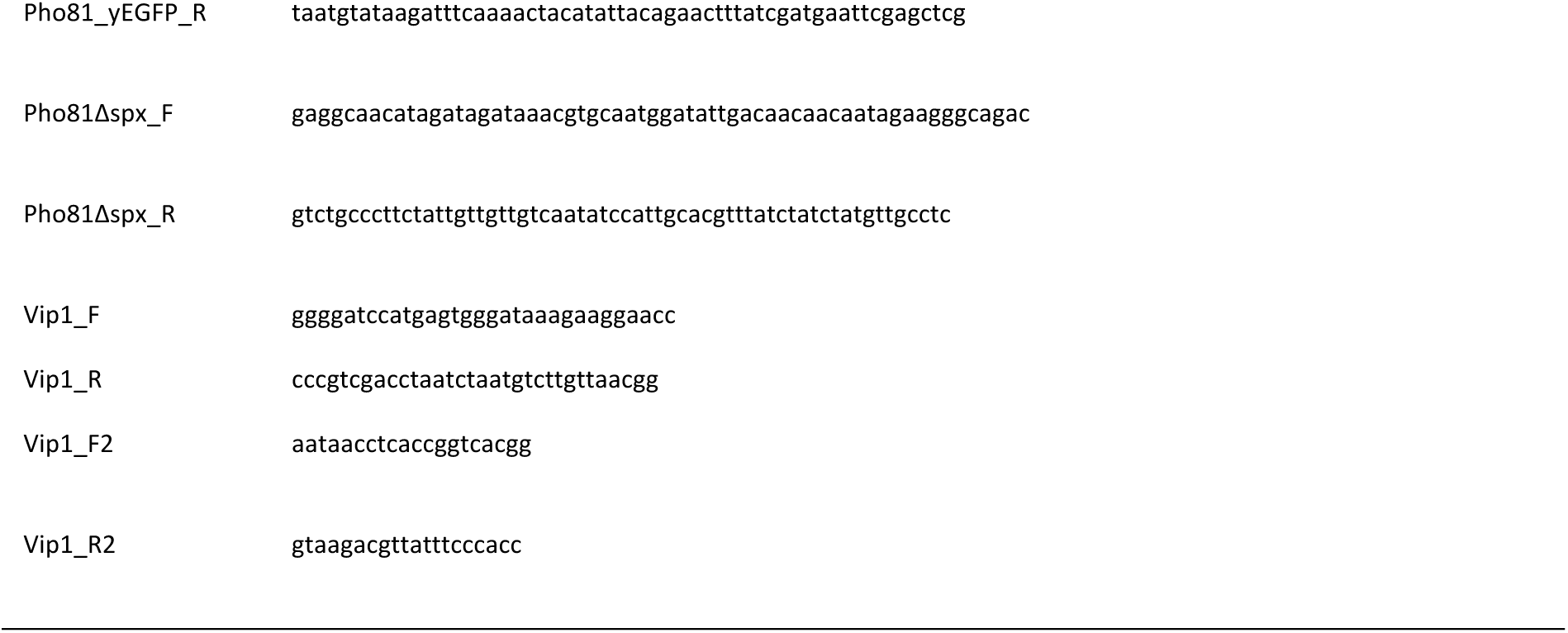
Oligonucleotides used in this study

### CE-ESI-MS parameters

All experiments were performed with a bare fused silica capillary with a length of 100 cm and 50 μm internal diameter. 35 mM ammonium acetate titrated by ammonia solution to pH 9.7 was used as background electrolyte (BGE). The sheath liquid was composed of a water-isopropanol (1:1) mixture, with a flow rate of 10 µL/min. 15 µL Inositol pyrophosphate extracts was spiked with 0.75 µL isotopic internal standards mixture (2 µM [^13^C_6_]1,5-IP_8_, 10 µM [^13^C_6_]5-IP_7_, 10 µM [^13^C_6_]1-IP_7_ and 40 µM [^13^C_6_]IP_6_). Samples were introduced by applying 100 mbar pressure for 10 s (20 nL). A separation voltage of +30 kV was applied over the capillary, generating a constant CE current at around 19 µA.

The MS source parameters setting were as follows: nebulizer pressure was 8 psi, sheath gas temperature was 175 °C and with a flow at 8 L/min, gas temperature was 150 °C and, with a flow of 11 L/min, the capillary voltage was -2000 V with nozzle voltage 2000V. Negative high pressure RF and low pressure RF (Ion Funnel parameters) was 70V and 40V, respectively. Parameters for MRM transitions are in Table S4.

**Table S4.** Parameters for MRM transitions

| Compound | Precursor Ion | Product Ion | dwell | CE (V) | Cell Acc (V) | Polarity |
| --- | --- | --- | --- | --- | --- | --- |
| [<sup>13</sup>C<sub>6</sub>]IP<sub>8</sub> | 411.9 | 362.9 | 80 | 10 | 1 | Negative |
| IP<sub>8</sub> | 408.9 | 359.9 | 80 | 10 | 1 | Negative |
| $[^{13}\text{C}_6]\text{IP}_7$ | 371.9 | 322.9 | 80 | 10 | 1 | Negative |
| $\text{IP}_7$ | 368.9 | 319.9 | 80 | 10 | 1 | Negative |
| $[^{13}\text{C}_6]\text{IP}_6$ | 331.9 | 486.9 | 80 | 17 | 1 | Negative |
| $\text{IP}_6$ | 328.9 | 480.9 | 80 | 17 | 1 | Negative |

## Supplementary Figures

**Fig. S1.**
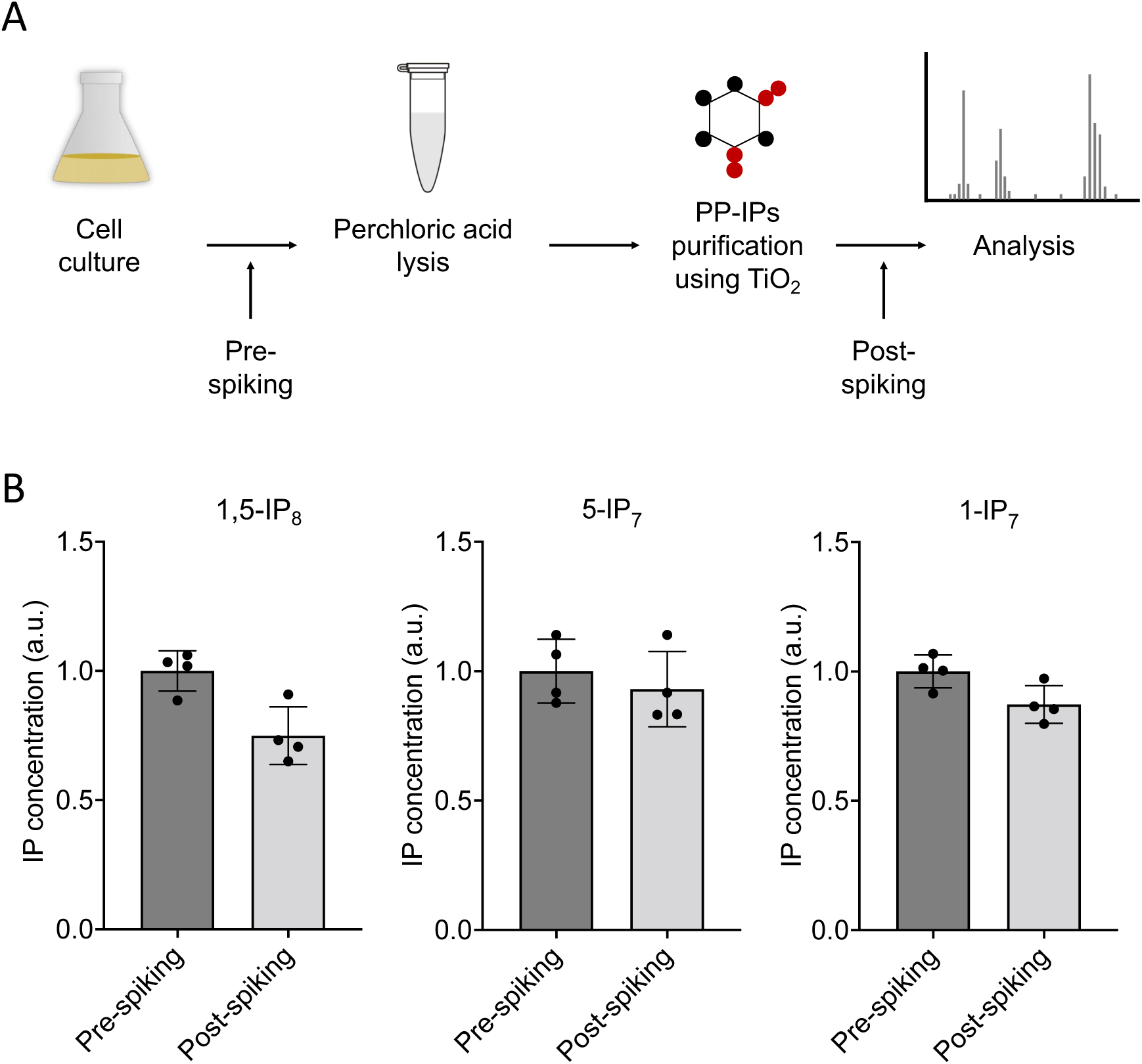
Recovery of IPPs extracted from *S. cerevisiae* cells. (A) Illustration of the procedure of IPP extraction using TiO_2_ beads. 0.1 µM [^13^C_6_]1,5-IP_8_, 0.5 µM [^13^C_6_]5-IP_7_ and 0.5 µM [^13^C_6_]1-IP_7_ were spiked into the yeast culture immediately before perchloric acid extraction (pre-spiking), or spiked into the IPP fraction extracted from the cells before the CE-MS measurement (post-spiking). (B) Recovery of IPPs in the assay. IPP purification showed 75% of recovery for 1,5-IP_8_, 93% for 5-IP_7_ and 87% for 1-IP_7_. Means (n=4) and standard deviations are indicated.

**Fig. S2.**
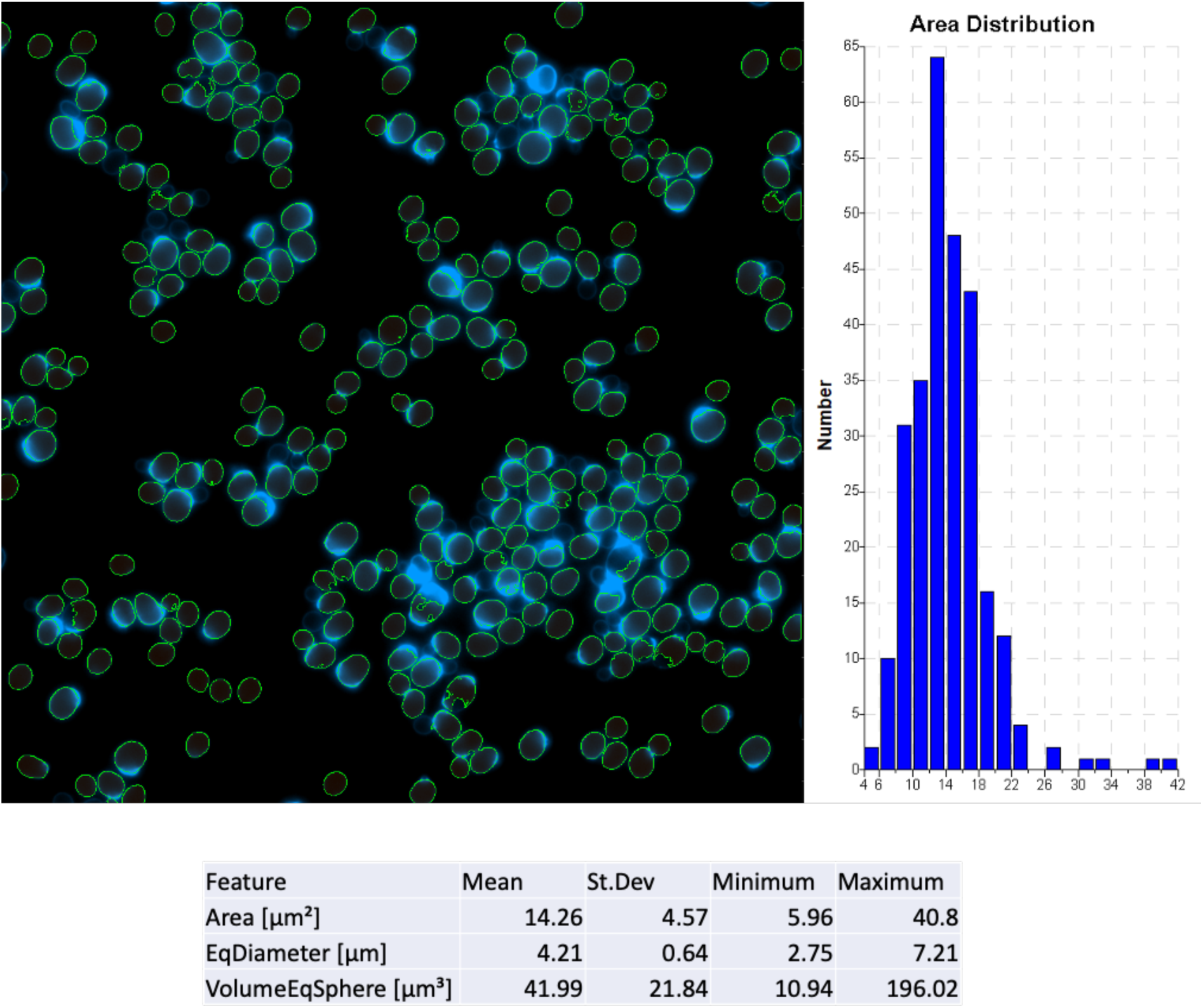
Determination of cell dimensions from wild-type cells stained with trypan blue. Cells were grown in SC medium over night until they reached an OD_600nm_ of 1. 10 µg/mL of Trypan Blue was added, and the cells were analyzed on a fluorescence microscope. Cells in the acquired images were analyzed by automated image segmentation and their volume was measured using the Nikon NIS Elements General Analysis 3 software package. The script for measuring cell volumes has been deposited at Figshare under the DOI <u>10.6084/m9.figshare.c.6700281.</u>

**Fig. S3:**
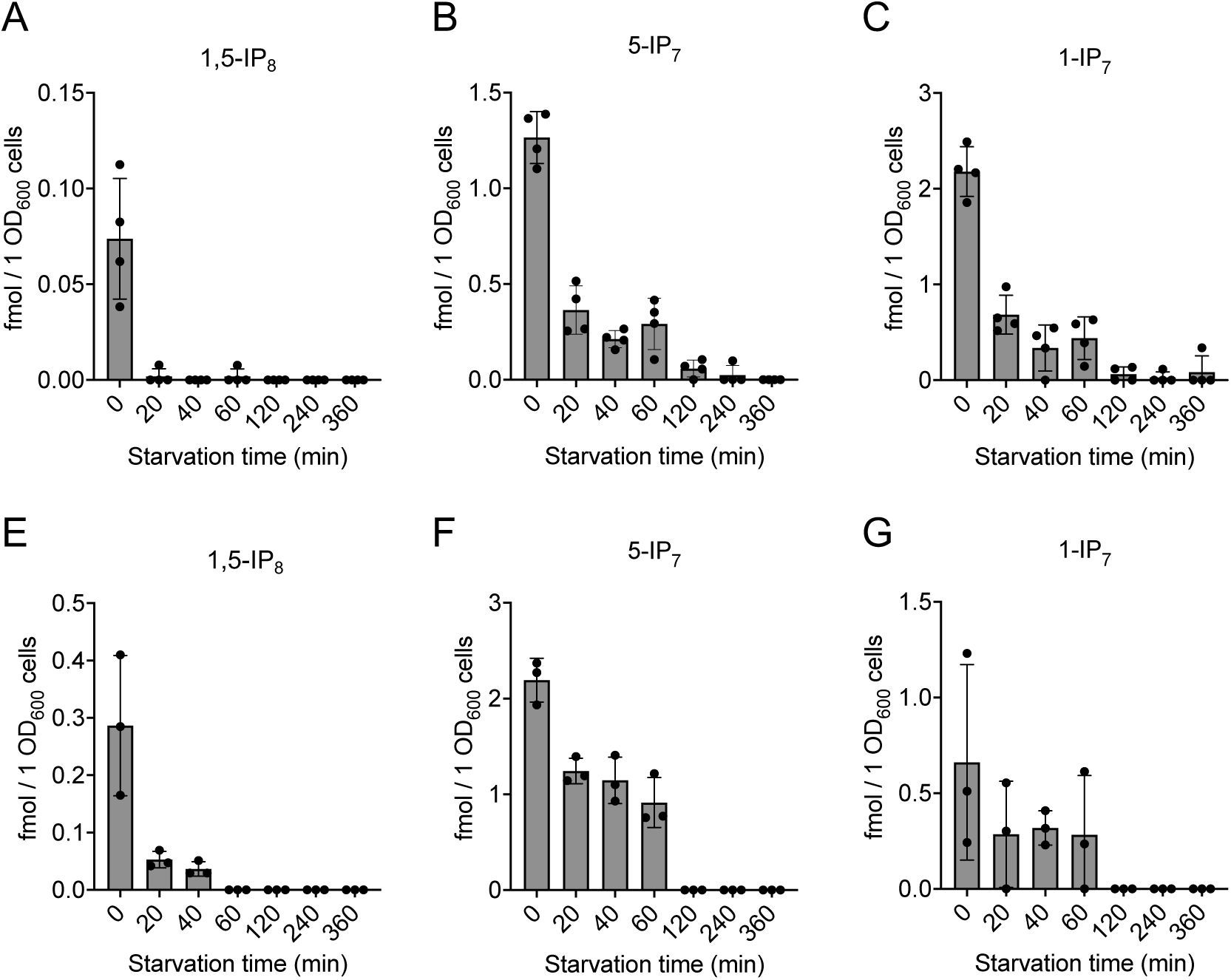
**Inositol pyrophosphate analysis in *C. neoformans* and *S. pombe*** Inositol pyrophosphates were measured in *C. neoformans* (A-C) and *S. pombe* (D-F). Both fungi were logarithmically grown in synthetic complete (SC) medium for 17 h up to an OD_600nm_ of 1. Cells were sedimented by centrifugation, resuspended in SC without P_i_, and incubated further. At the indicated times, aliquots were extracted with perchloric acid. IPPs were enriched on TiO_2_ beads and analyzed by CE-MS. Concentrations of 1,5-IP_8_, 5-IP_7_ and 1-IP_7_ in the extracts were determined by comparison with added synthetic ^13^C-labeled IPP standards. The graphs provide the concentrations in the extracts. n=4 for *C. neoformans*, and n=3 for *S. pombe*; means and standard deviations are indicated. The IPP values were normalized to the OD_600_ of the culture for every given time point.

**Fig. S4.**
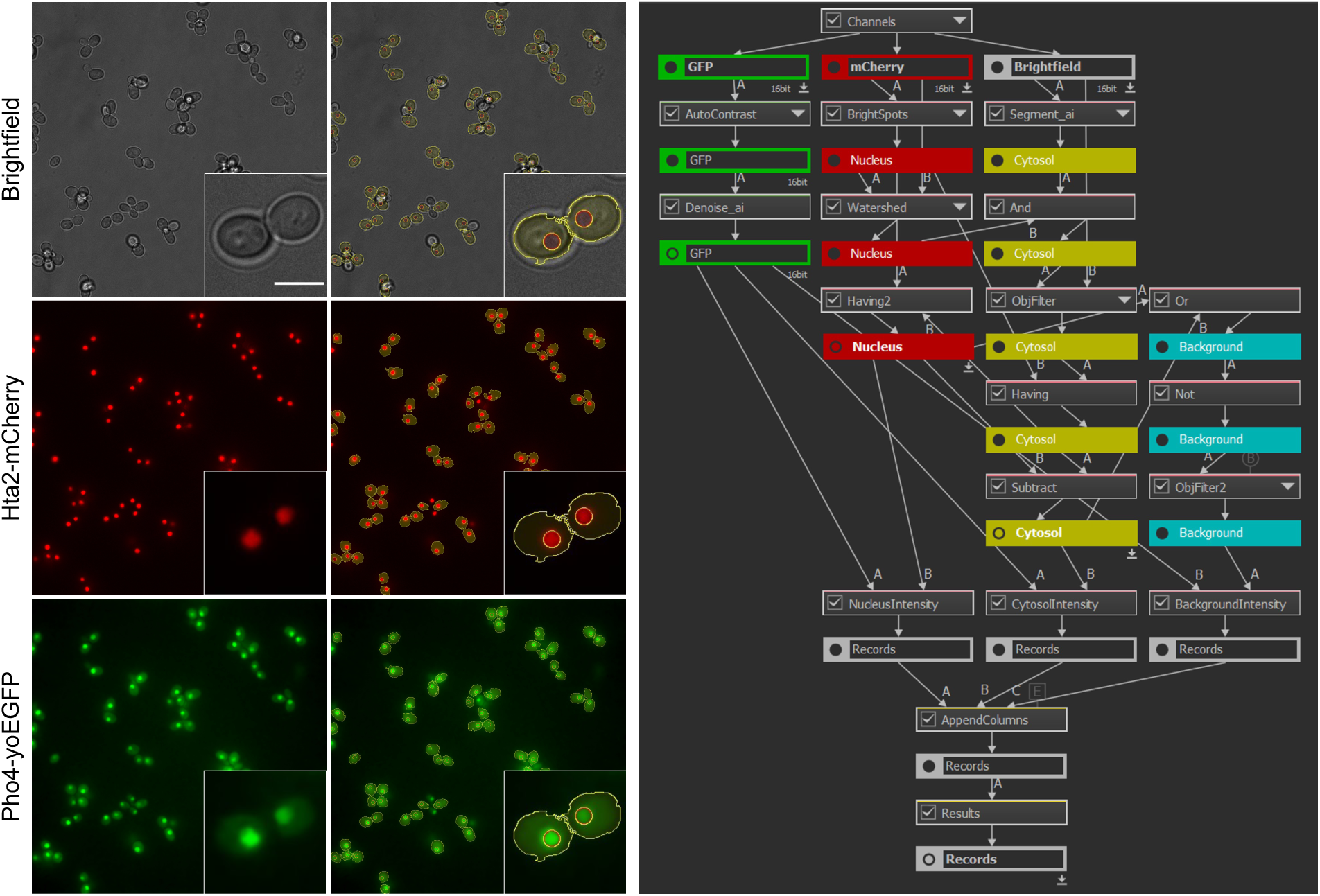
Segmentation of fluorescence microscopy time-lapse experiments. The subcellular localization of Pho4-yEGFP has been quantified from time-lapse fluorescence microscopy experiments. This quantification required the segmentation of microscopy images to discriminate cytosolic and nuclear compartments. To this end, the Hta2 histone of Pho4-yEGFP strains has been tagged with mCherry. Nuclei were segmented using mCherry fluorescence images while the cell contours were segmented using brightfield images. Cell segmentation and the fluorescence quantification were performed using the General Analysis 3 module of the NIS Elements software (Nikon). (A) Brightfield and two fluorescence images of the same field are shown. Yellow lines indicate the boundaries of the cells and nuclei that have been recognized by the algorithm. (B) Flow chart of the commands of the NIS Elements General Analysis 3 suite used for the segmentation. The script has been deposited at Figshare under the doi <u>10.6084/m9.figshare.c.6700281</u>.

**Fig. S5:**
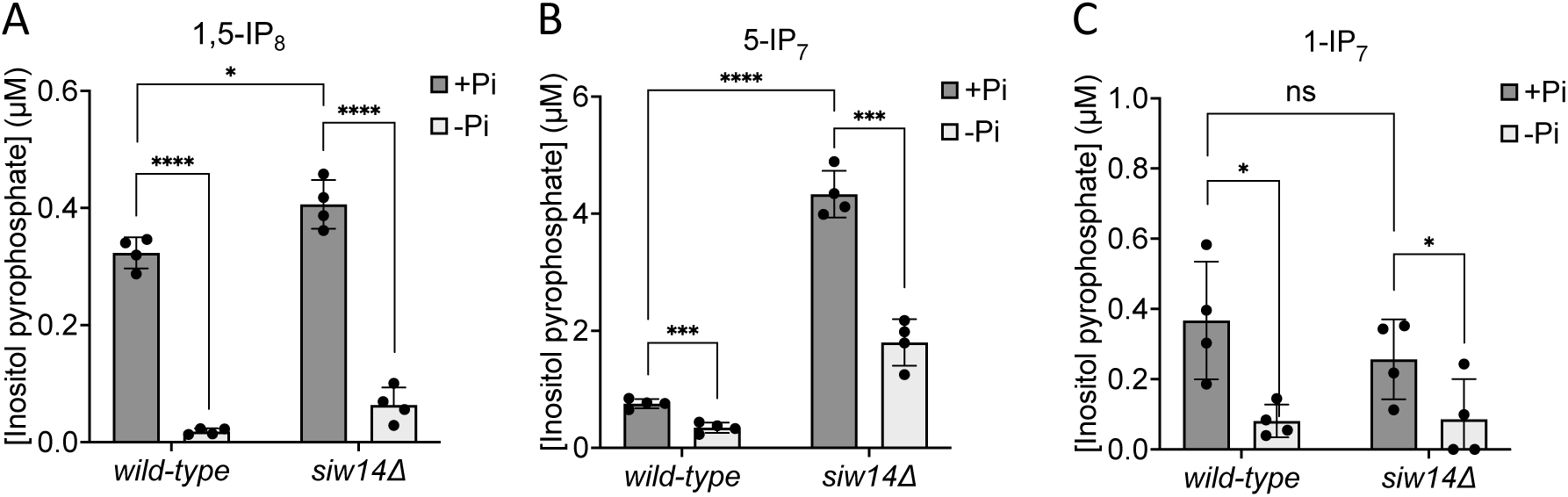
Loss of IPPs from *siw14Δ* cells upon P_i_ starvation. The indicated strains were grown logarithmically in SC medium containing 7.5 mM of P_i_ (30°C, 150 rpm, overnight). The cells were spun down, washed twice with P_i_ starvation medium and further incubated in P_i_ starvation medium or in SC with Pi for 2h. The inoculum for the samples for this final 2h incubation was adjusted such that all samples had an OD_600nm_ of 1 (10^7^ cells/ml) at the time of harvesting. After the two hours incubation, 1 ml of culture was extracted with perchloric acid and analyzed for IPPs by CE-ESI-MS. The data was normalized by the number of cells harvested before calculating cytosolic concentrations. The y-axis provides the estimated cytosolic concentrations based on an average cell volume of 42 fL. Means (n=4) and standard deviations are indicated. Graphs show the content of (A) 1,5-IP_8_, (B) 5-IP_7_, and (C) 1-IP_7._ **** p<0.0001; *** p<0.001; ** p<0.01; * p<0.05; n.s. not significant, determined with Student’s t-test.

**Fig. S6.**
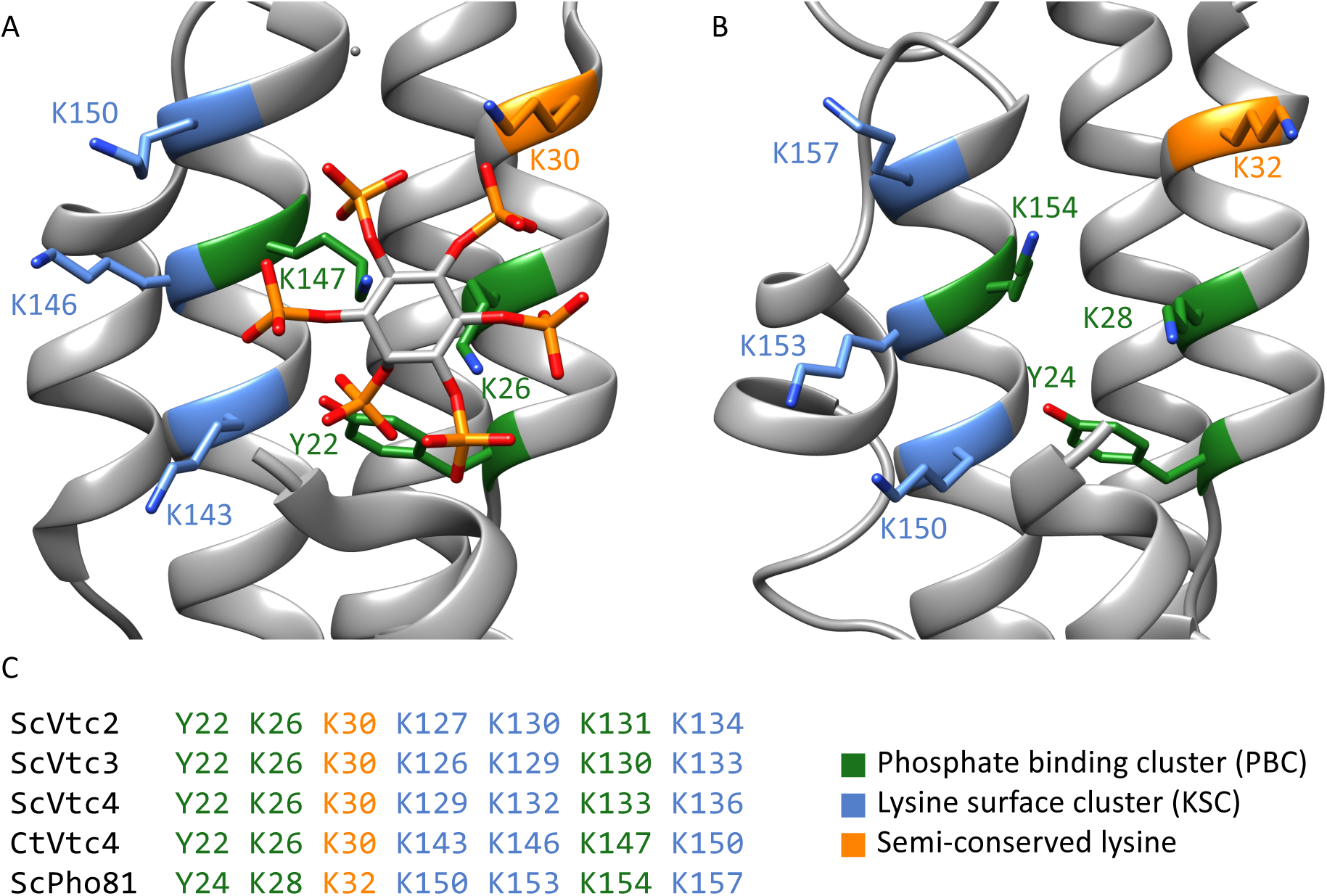
***Myo*-inositol polyphosphate binding pocket of yeast SPX domains.** (A) Crystal structure close-up view of *C. thermophilum* Vtc4 SPX domain complexing IP6 (5IJP). (B) Structure prediction (generated with SWISS-MODEL) of the *S. cerevisiae* Pho81 SPX domain revealing the potential conserved PBC and KSC clusters. (C) Alignment of the conserved residues *myo*-inositol polyphosphate binding motifs of Pho81 and of the different VTC subunits.

**Fig. S7:**
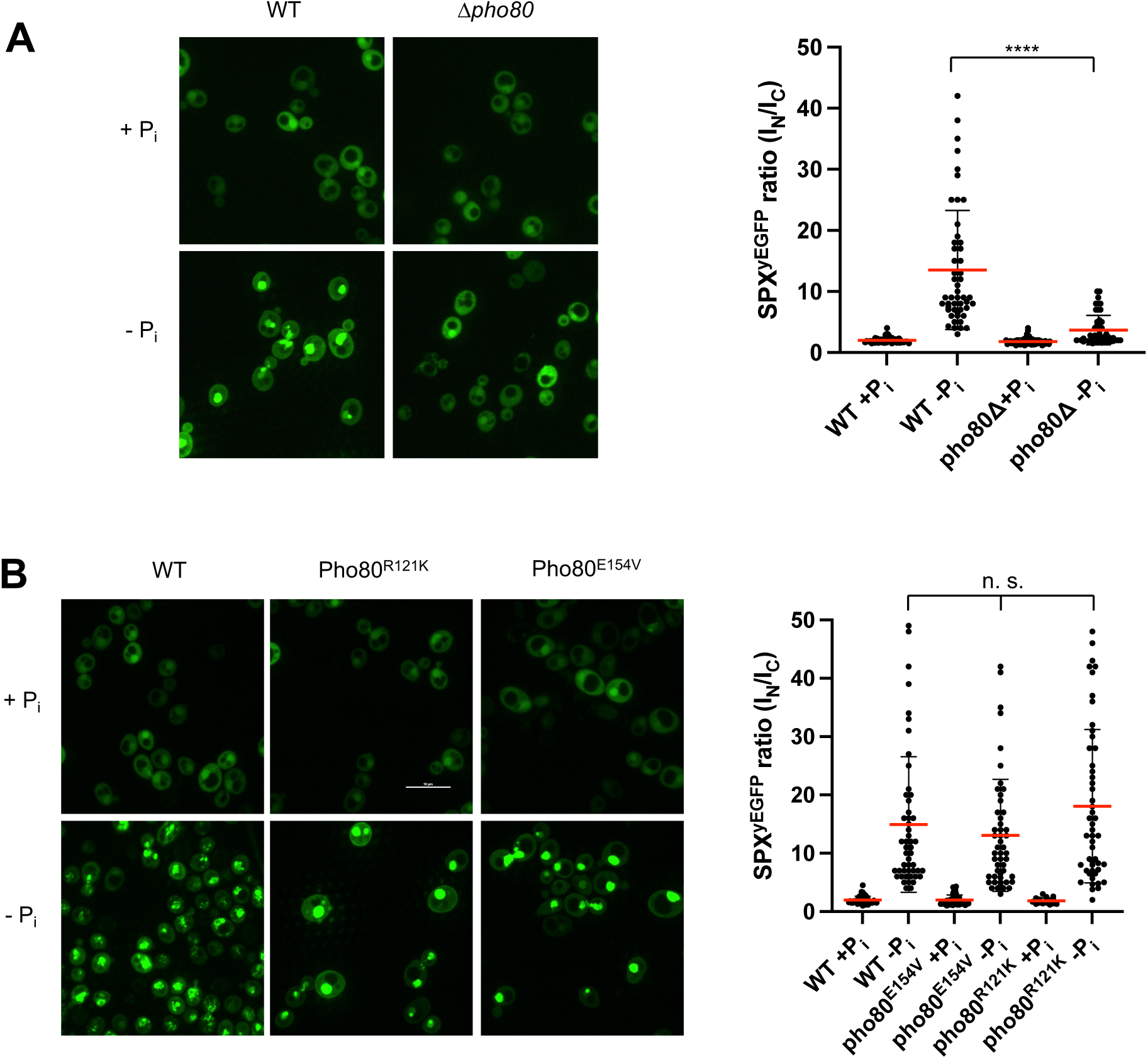
Recruitment of Pho81SPX^yEGFP^ to the nucleus by Pho85-Pho80. The indicated wildtype or isogenic mutant cells expressing the SPX domain of Pho81 as a yEGFP fusion (Pho81SPX^yEGFP^) from a centromeric plasmid under the ADH promotor were logarithmically grown in SC, sedimented in a tabletop centrifuge, and transferred into SC with (+P_i_) or without P_i_ (-P_i_). After 3 h of further cultivation, cells were imaged by spinning disc confocal microscopy. 50 cells per condition were analyzed from two independent experiments. Regions of interest were defined manually, and the fluorescence contained in the nuclei (I_N_) or the cytosol (I_C_) was integrated using ImageJ. Scale bar: 5 µm. Indicated pairwise differences were evaluated by a Mann-Whitney test. **** p<0.0001; n.s. not significant. (A) WT (BY4742) and isogenic *pho8011* cells expressing *PHO81SPX^yEGFP^*. (B) WT (BY4741) and isogenic cells expressing *pho80^R121K^* or *pho80^E154V^* from their genomic locus and Pho81SPX^yEGFP^ from the plasmid.

## References

1. S. Austin, A. Mayer, Phosphate Homeostasis - A Vital Metabolic Equilibrium Maintained Through the INPHORS Signaling Pathway. Front Microbiol 11, 1367 (2020).

2. C. Azevedo, A. Saiardi, Eukaryotic Phosphate Homeostasis: The Inositol Pyrophosphate Perspective. Trends Biochem Sci 42, 219–231 (2017).

3. D. Secco, et al., The emerging importance of the SPX domain-containing proteins in phosphate homeostasis. New Phytol 193, 842–851 (2012).

4. R. Gerasimaite, et al., Inositol Pyrophosphate Specificity of the SPX-Dependent Polyphosphate Polymerase VTC. ACS Chem. Biol. 12, 648–653 (2017).

5. E. Potapenko, et al., 5-Diphosphoinositol Pentakisphosphate (5-IP7) Regulates Phosphate Release from Acidocalcisomes and Yeast Vacuoles. J. Biol. Chem., jbc.RA118.005884 (2018).

6. R. Wild, et al., Control of eukaryotic phosphate homeostasis by inositol polyphosphate sensor domains. Science 352, 986–990 (2016).

7. M. K. Ried, et al., Inositol pyrophosphates promote the interaction of SPX domains with the coiled-coil motif of PHR transcription factors to regulate plant phosphate homeostasis. Nature communications 12, 1–13 (2021).

8. C. Wang, et al., Rice SPX-Major Facility Superfamily3, a Vacuolar Phosphate Efflux Transporter, Is Involved in Maintaining Phosphate Homeostasis in Rice. Plant Physiol 169, 2822–2831 (2015).

9. T.-Y. Liu, et al., Identification of plant vacuolar transporters mediating phosphate storage. Nature communications 7, 11095 (2016).

10. J. Dong, et al., Inositol Pyrophosphate InsP8 Acts as an Intracellular Phosphate Signal in Arabidopsis. Mol Plant 12, 1463–1473 (2019).

11. W. Liu, et al., Cryo-EM structure of the polyphosphate polymerase VTC reveals coupling of polymer synthesis to membrane transit. Embo J (2023) https://doi.org/10.15252/embj.2022113320.

12. J. Pipercevic, et al., Inositol pyrophosphates activate the vacuolar transport chaperone complex in yeast by disrupting a homotypic SPX domain interaction. Nat Commun 14, 2645 (2023).

13. Z. Guan, et al., The cytoplasmic synthesis and coupled membrane translocation of eukaryotic polyphosphate by signal-activated VTC complex. Nat Commun 14, 718 (2023).

14. X. Li, et al., Control of XPR1-dependent cellular phosphate efflux by InsP8 is an exemplar for functionally-exclusive inositol pyrophosphate signaling. Proc. Natl. Acad. Sci. U.S.A. 14, 201908830 (2020).

15. M. S. Wilson, H. J. Jessen, A. Saiardi, The inositol hexakisphosphate kinases IP6K1 and -2 regulate human cellular phosphate homeostasis, including XPR1-mediated phosphate export. J. Biol. Chem., jbc.RA119.007848 (2019).

16. E. Riemer, et al., ITPK1 is an InsP6/ADP phosphotransferase that controls phosphate signaling in Arabidopsis. Mol Plant (2021) https://doi.org/10.1016/j.molp.2021.07.011.

17. D. Laha, et al., VIH2 Regulates the Synthesis of Inositol Pyrophosphate InsP8 and Jasmonate-Dependent Defenses in Arabidopsis. Plant Cell 27, 1082–1097 (2015).

18. J. Zhu, et al., Two bifunctional inositol pyrophosphate kinases/phosphatases control plant phosphate homeostasis. elife 8 (2019).

19. Y. S. Lee, S. Mulugu, J. D. York, E. K. O’Shea, Regulation of a Cyclin-CDK-CDK Inhibitor Complex by Inositol Pyrophosphates. Science 316, 109–112 (2007).

20. Y.-S. Lee, K. Huang, F. A. Quiocho, E. K. O’Shea, Molecular basis of cyclin-CDK-CKI regulation by reversible binding of an inositol pyrophosphate. Nat Chem Biol 4, 25–32 (2008).

21. D. Desmarini, et al., IP7-SPX Domain Interaction Controls Fungal Virulence by Stabilizing Phosphate Signaling Machinery. MBio 11, 873 (2020).

22. A. Lonetti, et al., Identification of an evolutionary conserved family of inorganic polyphosphate endopolyphosphatases. J Biol Chem (2011) https://doi.org/10.1074/jbc.m111.266320.

23. C. Auesukaree, H. Tochio, M. Shirakawa, Y. Kaneko, S. Harashima, Plc1p, Arg82p, and Kcs1p, enzymes involved in inositol pyrophosphate synthesis, are essential for phosphate regulation and polyphosphate accumulation in Saccharomyces cerevisiae. J Biol Chem 280, 25127–25133 (2005).

24. M. Pascual-Ortiz, E. Walla, U. Fleig, A. Saiardi, The PPIP5K Family Member Asp1 Controls Inorganic Polyphosphate Metabolism in S. pombe. J Fungi (Basel) 7, 626 (2021).

25. H. Wang, et al., Asp1 from Schizosaccharomyces pombe binds a [2Fe-2S](2+) cluster which inhibits inositol pyrophosphate 1-phosphatase activity. Biochemistry 54, 6462–6474 (2015).

26. T. A. Randall, C. Gu, X. Li, H. Wang, S. B. Shears, A two-way switch for inositol pyrophosphate signaling: Evolutionary history and biological significance of a unique, bifunctional kinase/phosphatase. Adv Biol Regul, 100674 (2019).

27. M. Pascual-Ortiz, et al., Asp1 Bifunctional Activity Modulates Spindle Function via Controlling Cellular Inositol Pyrophosphate Levels in Schizosaccharomyces pombe. Mol Cell Biol 38, MCB.00047–18 (2018).

28. B. Topolski, V. Jakopec, N. A. Künzel, U. Fleig, The inositol pyrophosphate kinase Asp1 modulates chromosome segregation fidelity and spindle function in S. pombe. *Mol Cell Biol*, MCB.00330–16 (2016).

29. J. Pöhlmann, et al., The Vip1 inositol polyphosphate kinase family regulates polarized growth and modulates the microtubule cytoskeleton in fungi. PLoS Genet 10, e1004586 (2014).

30. A. M. Sanchez, A. Garg, S. Shuman, B. Schwer, Inositol pyrophosphates impact phosphate homeostasis via modulation of RNA 3’ processing and transcription termination. Nucleic Acids Res 47, 8452–8469 (2019).

31. B. Benjamin, et al., Activities and Structure-Function Analysis of Fission Yeast Inositol Pyrophosphate (IPP) Kinase-Pyrophosphatase Asp1 and Its Impact on Regulation of pho1 Gene Expression. Mbio, e01034–22 (2022).

32. M. Estill, C. L. Kerwin-Iosue, D. D. Wykoff, Dissection of the PHO pathway in Schizosaccharomyces pombe using epistasis and the alternate repressor adenine. Curr Genet 61, 175–183 (2015).

33. T. C. Henry, et al., Systematic screen of Schizosaccharomyces pombe deletion collection uncovers parallel evolution of the phosphate signal transduction pathway in yeasts. Eukaryotic Cell 10, 198–206 (2011).

34. I. Carter-O’Connell, M. T. Peel, D. D. Wykoff, E. K. O’Shea, Genome-wide characterization of the phosphate starvation response in Schizosaccharomyces pombe. BMC Genomics 13, 697 (2012).

35. B. Schwer, A. Garg, A. Jacewicz, S. Shuman, Genetic screen for suppression of transcriptional interference identifies a gain-of-function mutation in Pol2 termination factor Seb1. Proc. Natl. Acad. Sci. U.S.A. 118 (2021).

36. A. M. Sanchez, S. Shuman, B. Schwer, RNA polymerase II CTD interactome with 3ʹ processing and termination factors in fission yeast and its impact on phosphate homeostasis. Proc National Acad Sci 115, E10652–E10661 (2018).

37. B. Schwer, A. M. Sanchez, S. Shuman, RNA polymerase II CTD phospho-sites Ser5 and Ser7 govern phosphate homeostasis in fission yeast. Rna New York N Y 21, 1770–80 (2015).

38. B. Schwer, D. A. Bitton, A. M. Sanchez, J. Bähler, S. Shuman, Individual letters of the RNA polymerase II CTD code govern distinct gene expression programs in fission yeast. Proc National Acad Sci 111, 4185–4190 (2014).

39. M. S. C. Wilson, A. Saiardi, Importance of Radioactive Labelling to Elucidate Inositol Polyphosphate Signalling. Top Curr Chem (Cham) 375, 14 (2017).

40. E. Eskes, M.-A. Deprez, T. Wilms, J. Winderickx, pH homeostasis in yeast; the phosphate perspective. Curr Genet 274, 30052–7 (2017).

41. M. Conrad, et al., Nutrient sensing and signaling in the yeast Saccharomyces cerevisiae. FEMS Microbiol Rev 38, 254–299 (2014).

42. P. Korber, S. Barbaric, The yeast PHO5 promoter: from single locus to systems biology of a paradigm for gene regulation through chromatin. Nucleic Acids Res 42, 10888–10902 (2014).

43. K. R. Schneider, R. L. Smith, E. K. O’Shea, Phosphate-regulated inactivation of the kinase PHO80-PHO85 by the CDK inhibitor PHO81. Science 266, 122–126 (1994).

44. N. Ogawa, et al., Functional domains of Pho81p, an inhibitor of Pho85p protein kinase, in the transduction pathway of Pi signals in Saccharomyces cerevisiae. Mol Cell Biol 15, 997– 1004 (1995).

45. K. Yoshida, N. Ogawa, Y. Oshima, Function of the PHO regulatory genes for repressible acid phosphatase synthesis in Saccharomyces cerevisiae. Mol. Gen. Genet. 217, 40–46 (1989).

46. A. Kaffman, N. M. Rank, E. K. O’Shea, Phosphorylation regulates association of the transcription factor Pho4 with its import receptor Pse1/Kap121. Genes Dev 12, 2673–2683 (1998).

47. A. Kaffman, N. M. Rank, E. M. O’Neill, L. S. Huang, E. K. O’Shea, The receptor Msn5 exports the phosphorylated transcription factor Pho4 out of the nucleus. Nature 396, 482– 486 (1998).

48. A. Komeili, E. K. O’Shea, Roles of phosphorylation sites in regulating activity of the transcription factor Pho4. Science 284, 977–980 (1999).

49. S. Huang, D. A. Jeffery, M. D. Anthony, E. K. O’Shea, Functional analysis of the cyclin-dependent kinase inhibitor Pho81 identifies a novel inhibitory domain. Mol Cell Biol 21, 6695–6705 (2001).

50. B. H. Spain, D. Koo, M. Ramakrishnan, B. Dzudzor, J. Colicelli, Truncated forms of a novel yeast protein suppress the lethality of a G protein alpha subunit deficiency by interacting with the beta subunit. J Biol Chem 270, 25435–25444 (1995).

51. A. Toh-E, Y. Oshima, Characterization of a dominant, constitutive mutation, PHOO, for the repressible acid phosphatase synthesis in Saccharomyces cerevisiae. J Bacteriol 120, 608–17 (1974).

52. C. L. Creasy, S. L. Madden, L. W. Bergman, Molecular analysis of the PHO81 gene of Saccharomyces cerevisiae. Nucleic Acids Res 21, 1975–1982 (1993).

53. C. L. Creasy, D. Shao, L. W. Begman, Negative transcriptional regulation of PH081 expression in Saccharomyces cerevisiae. Gene 168, 23–29 (1996).

54. G.-D. Kim, D. Qiu, H. J. Jessen, A. Mayer, Metabolic Consequences of Polyphosphate Synthesis and Imminent Phosphate Limitation. Mbio, e00102–23 (2023).

55. D. Qiu, et al., Analysis of inositol phosphate metabolism by capillary electrophoresis electrospray ionization mass spectrometry. Nature communications 11, 6035–12 (2020).

56. D. Qiu, et al., Capillary electrophoresis mass spectrometry identifies new isomers of inositol pyrophosphates in mammalian tissues. Chem Sci 14, 658–667 (2022).

57. S. Mulugu, et al., A conserved family of enzymes that phosphorylate inositol hexakisphosphate. Science 316, 106–109 (2007).

58. A. Saiardi, H. Erdjument-Bromage, A. M. Snowman, P. Tempst, S. H. Snyder, Synthesis of diphosphoinositol pentakisphosphate by a newly identified family of higher inositol polyphosphate kinases. Curr Biol 9, 1323–1326 (1999).

59. G. Zong, S. B. Shears, H. Wang, Structural and catalytic analyses of the InsP6 kinase activities of higher plant ITPKs. Faseb J 36, e22380 (2022).

60. H. Wang, J. R. Falck, T. M. T. Hall, S. B. Shears, Structural basis for an inositol pyrophosphate kinase surmounting phosphate crowding. Nat Chem Biol 8, 111–116 (2012).

61. G. Zong, et al., New structural insights reveal an expanded reaction cycle for inositol pyrophosphate hydrolysis by human DIPP1. FASEB J 35, e21275 (2021).

62. H. Wang, C. Gu, R. J. Rolfes, H. J. Jessen, S. B. Shears, Structural and biochemical characterization of Siw14: A protein-tyrosine phosphatase fold that metabolizes inositol pyrophosphates. J. Biol. Chem. 293, 6905–6914 (2018).

63. E. A. Steidle, et al., The InsP7 phosphatase Siw14 regulates inositol pyrophosphate levels to control localization of the general stress response transcription factor Msn2. J. Biol. Chem., jbc.RA119.012148 (2019).

64. D. E. Dollins, et al., Vip1 is a kinase and pyrophosphatase switch that regulates inositol diphosphate signaling. Proc. Natl. Acad. Sci. U.S.A. 117, 9356–9364 (2020).

65. R. K. Harmel, et al., Harnessing 13C-labeled myo-inositol to interrogate inositol phosphate messengers by NMR. Chem Sci 10, 5267–5274 (2019).

66. R. Puschmann, R. K. Harmel, D. Fiedler, Scalable Chemoenzymatic Synthesis of Inositol Pyrophosphates. Biochemistry 58, 3927–3932 (2019).

67. M. Uchida, et al., Quantitative analysis of yeast internal architecture using soft X-ray tomography. Yeast 28, 227–236 (2011).

68. D. Laha, et al., Arabidopsis ITPK1 and ITPK2 Have an Evolutionarily Conserved Phytic Acid Kinase Activity. ACS Chem. Biol. 14, 2127–2133 (2019).

69. O. Adepoju, et al., Inositol Trisphosphate Kinase and Diphosphoinositol Pentakisphosphate Kinase Enzymes Constitute the Inositol Pyrophosphate Synthesis Pathway in Plants. bioRxiv, 724914 (2019).

70. H. Whitfield, et al., An ATP-responsive metabolic cassette comprised of inositol tris/tetrakisphosphate kinase 1 (ITPK1) and inositol pentakisphosphate 2-kinase (IPK1) buffers diphosphosphoinositol phosphate levels. Biochem J 477, 2621–2638 (2020).

71. M. N. Trung, et al., Stable Isotopomers of myo-Inositol Uncover a Complex MINPP1-Dependent Inositol Phosphate Network. Acs Central Sci 8, 1683–1694 (2022).

72. C. Azevedo, A. Saiardi, Extraction and analysis of soluble inositol polyphosphates from yeast. Nat Protoc 1, 2416–2422 (2006).

73. M. R. Thomas, E. K. O’Shea, An intracellular phosphate buffer filters transient fluctuations in extracellular phosphate levels. Proc Natl Acad Sci USA 102, 9565–9570 (2005).

74. D. D. Wykoff, A. H. Rizvi, J. M. Raser, B. Margolin, E. K. O’Shea, Positive feedback regulates switching of phosphate transporters in S. cerevisiae. Mol Cell 27, 1005–1013 (2007).

75. E. M. O’Neill, A. Kaffman, E. R. Jolly, E. K. O’Shea, Regulation of PHO4 nuclear localization by the PHO80-PHO85 cyclin-CDK complex. Science 271, 209–212 (1996).

76. M. Nishizawa, et al., Nutrient-regulated antisense and intragenic RNAs modulate a signal transduction pathway in yeast. PLoS Biol 6, 2817–2830 (2008).

77. A. Almer, H. Rudolph, A. Hinnen, W. Hörz, Removal of positioned nucleosomes from the yeast PHO5 promoter upon PHO5 induction releases additional upstream activating DNA elements. Embo J 5, 2689–2696 (1986).

78. S. Barbaric, et al., Redundancy of chromatin remodeling pathways for the induction of the yeast PHO5 promoter in vivo. Journal of Biological Chemistry 282, 27610–27621 (2007).

79. F. H. Lam, D. J. Steger, E. K. O’Shea, Chromatin decouples promoter threshold from dynamic range. Nature 453, 246–250 (2008).

80. B. Pinson, et al., Metabolic intermediates selectively stimulate transcription factor interaction and modulate phosphate and purine pathways. Genes Dev 23, 1399–1407 (2009).

81. M. Varadi, et al., AlphaFold Protein Structure Database: massively expanding the structural coverage of protein-sequence space with high-accuracy models. Nucleic Acids Res. 50, D439–D444 (2021).

82. M. Mirdita, et al., ColabFold: making protein folding accessible to all. Nat. Methods 19, 679–682 (2022).

83. K. Huang, et al., Structure of the Pho85-Pho80 CDK-cyclin complex of the phosphate-responsive signal transduction pathway. Mol Cell 28, 614–623 (2007).

84. M. R. Wolff, A. Schmid, P. Korber, U. Gerland, Effective dynamics of nucleosome configurations at the yeast PHO5 promoter. Elife 10, e58394 (2021).

85. M. Hothorn, et al., Catalytic core of a membrane-associated eukaryotic polyphosphate polymerase. Science 324, 513–516 (2009).

86. R. Gerasimaite, A. Mayer, Enzymes of yeast polyphosphate metabolism: structure, enzymology and biological roles. Biochem Soc Trans 44, 234–239 (2016).

87. B. Schwer, et al., Cleavage-Polyadenylation Factor Cft1 and SPX Domain Proteins Are Agents of Inositol Pyrophosphate Toxicosis in Fission Yeast. Mbio 13, e03476–21 (2022).

88. A. Garg, S. Shuman, B. Schwer, A genetic screen for suppressors of hyper-repression of the fission yeast PHO regulon by Pol2 CTD mutation T4A implicates inositol 1-pyrophosphates as agonists of precocious lncRNA transcription termination. Nucleic Acids Res 48, gkaa776-(2020).

89. A. Toh-e, et al., Identification of genes involved in the phosphate metabolism in Cryptococcus neoformans. Fungal Genet Biol 80, 19–30 (2015).

90. J. Choi, A. Rajagopal, Y.-F. Xu, J. D. Rabinowitz, E. K. O’Shea, A systematic genetic screen for genes involved in sensing inorganic phosphate availability in Saccharomyces cerevisiae. PLoS ONE 12, e0176085 (2017).

91. D. Giovannini, J. Touhami, P. Charnet, M. Sitbon, J.-L. Battini, Inorganic phosphate export by the retrovirus receptor XPR1 in metazoans. Cell Rep 3, 1866–1873 (2013).

92. M. A. Sheff, K. S. Thorn, Optimized cassettes for fluorescent protein tagging in Saccharomyces cerevisiae. Yeast 21, 661–670 (2004).

93. D. Mumberg, R. Müller, M. Funk, Yeast vectors for the controlled expression of heterologous proteins in different genetic backgrounds. Gene 156, 119–122 (1995).

